# Random auditory stimulation disturbs traveling slow waves and declarative memory

**DOI:** 10.1101/2025.02.06.636865

**Authors:** Nora M. Roüast, Deniz Kumral, Steffen Gais, Monika Schönauer

## Abstract

Sleep plays a crucial role in memory consolidation, and various methods have attempted to enhance this process by using auditory stimulation to modulate slow waves or trigger memory reactivation. However, the broader impact of auditory stimulation on sleep physiology and memory retention is not fully understood. Here, we apply random or sham auditory stimulation during an afternoon nap in a within-subject design to investigate its effects on declarative and procedural memory consolidation and electrical brain activity during sleep. Stimulation led to a specific reduction in slow-wave sleep, a decreased slow-wave count, and impaired declarative recall that correlated with the diminished sleep depth. Furthermore, we observed altered slow-wave traveling dynamics, with stimulation resulting in shorter traveling trajectories with less reach and reduced spatial spread, particularly impacting frontal regions. Disruptions in these features correlated with memory deficits. Our findings highlight the role of dynamic slow-wave properties in declarative memory consolidation and reveal some methodological limitations for using auditory cues during sleep, given their potential to disrupt complex processing patterns.

## Introduction

The dependence of memory consolidation (Rasch & Born, 2013) on sleep has long been established for procedural learning (Schönauer et al., 2013; Stickgold & Walker, 2005), but seems particularly strong for declarative memory processing (Diekelmann & Born, 2010; Plihal & Born, 1997; Schönauer et al., 2014). According to contemporary models and backed by a growing body of research (e.g., Belal et al., 2018; Ngo et al., 2020; Schönauer et al., 2017; Schreiner et al., 2023, 2024; Schreiner & Staudigl, 2020; Zhang et al., 2018), covert reactivation of mnemonic content underlies sleep-dependent memory consolidation by propagating information from hippocampal to neocortical regions via a synchronized interplay of hippocampal sharp-wave ripples, thalamic spindles, and neocortical slow waves (Born & Wilhelm, 2012; Klinzing et al., 2019; Staresina et al., 2015). In line with this idea, measures of spindle activity have been related to both declarative and procedural memory consolidation (Cairney et al., 2018; Gais et al., 2002; Kaestner et al., 2013; Kumral et al., 2023; Manoach et al., 2010; Nishida & Walker, 2007; Petzka et al., 2022; Schabus et al., 2006; Wamsley et al., 2012), and slow waves to the consolidation of declarative memory performance (Marshall et al., 2006; Ngo et al., 2013). Indeed, reactivation drives memory retention (Dupret et al., 2010) and interruption of the key oscillatory mechanisms thought to support this process impairs memory performance (Latchoumane et al., 2017; Wilckens et al., 2018). Therefore, the relationship between memory reactivation and sleep physiology presents a potential avenue for improving memory consolidation.

Targeted memory reactivation has successfully utilized auditory or olfactory cues during sleep to improve later memory retention in humans (Hu et al., 2020; Rasch et al., 2007). By presenting mnemonic cues linked to previous learning material, the re-emergence of specific learning-related activity patterns can be triggered during sleep such that this information gets preferentially consolidated. Moreover, the general opportunity for memory consolidation could be increased by boosting the supportive physiology, such as slow waves. Both transcranial magnetic and auditory stimulation were used to trigger or enhance slow waves (Bergmann et al., 2012; Besedovsky et al., 2017; Esfahani et al., 2023; Massimini et al., 2007; Ngo et al., 2013). The timing of this auditory stimulation appears to be crucial, as cues synchronized with the slow-wave upstate enhance slow wave activity and may consequently improve memory retention (Besedovsky et al., 2017; Esfahani et al., 2023; Ngo et al., 2013; but see Henin et al., 2019; Wunderlin et al., 2021). Cues applied during identified slow waves irrespective of phase may lead to their suppression, even though the effects on memory are still unclear (Fehér et al., 2023). Importantly though, noise generally has been associated with detrimental effects of sleep depth, even if the overall duration remains unchanged (Fehér et al., 2023; Muzet, 2007; Schönauer et al., 2014). Thus, auditory input during sleep can both lead to functional gains or losses, with the ultimate memory benefit of sleep potentially being the sum of both influences and depending on ties to content or timing.

Here, we aimed to investigate how randomly occurring sounds, unrelated to prior learning material and not synchronized with the slow wave phase, influence sleep-dependent memory consolidation and alter sleep-physiological processes crucial for memory processing. Could randomly administered auditory stimulation be a mechanism to disrupt deep sleep and consequently mnemonic processing? Additionally, sounds can alter the dynamic propagation of slow waves (Sousouri et al., 2022). Yet, the traveling behavior of slow waves may depend on undisturbed sleep, allowing specific states of brain connectivity to emerge which serve memory processing (Avvenuti et al., 2020; Castelnovo et al., 2023; Kurth et al., 2017; Massimini et al., 2004; Schoch et al., 2018; Sharon et al., 2024). Disturbing traveling behavior and therefore natural information propagation across the brain might consequently deteriorate memory consolidation. To examine the effect of random auditory stimulation on memory processing during sleep, we conducted a within-subject EEG study with 20 healthy young adults. Participants studied both procedural and declarative learning tasks before a nap opportunity that once included random auditory stimulation and once sham stimulation (silence). Auditory stimulation during sleep led to a marked decrease in slow wave sleep (SWS) and slow oscillatory (SO) activity, which was specifically associated with impaired declarative memory. Furthermore, stimulation stunted the traveling profiles of slow waves, with affected markers significantly correlating with declarative memory deficits.

## Results

Twenty healthy participants (age: 24.1±3.9; 20 male) came for two experimental sessions, during which they encoded declarative and procedural memory content before a three-hour nap opportunity, and retrieved the information subsequently (Fig. 1a, see Methods). In the stimulation session, sounds were played throughout the nap opportunity, whilst in the sham session no sounds were presented. Electroencephalography (EEG) recordings during the nap period as well as retrieval performance served as markers of the effect of stimulation on sleep quality and physiology as well as the potential behavioral cost.

**Figure 1.**
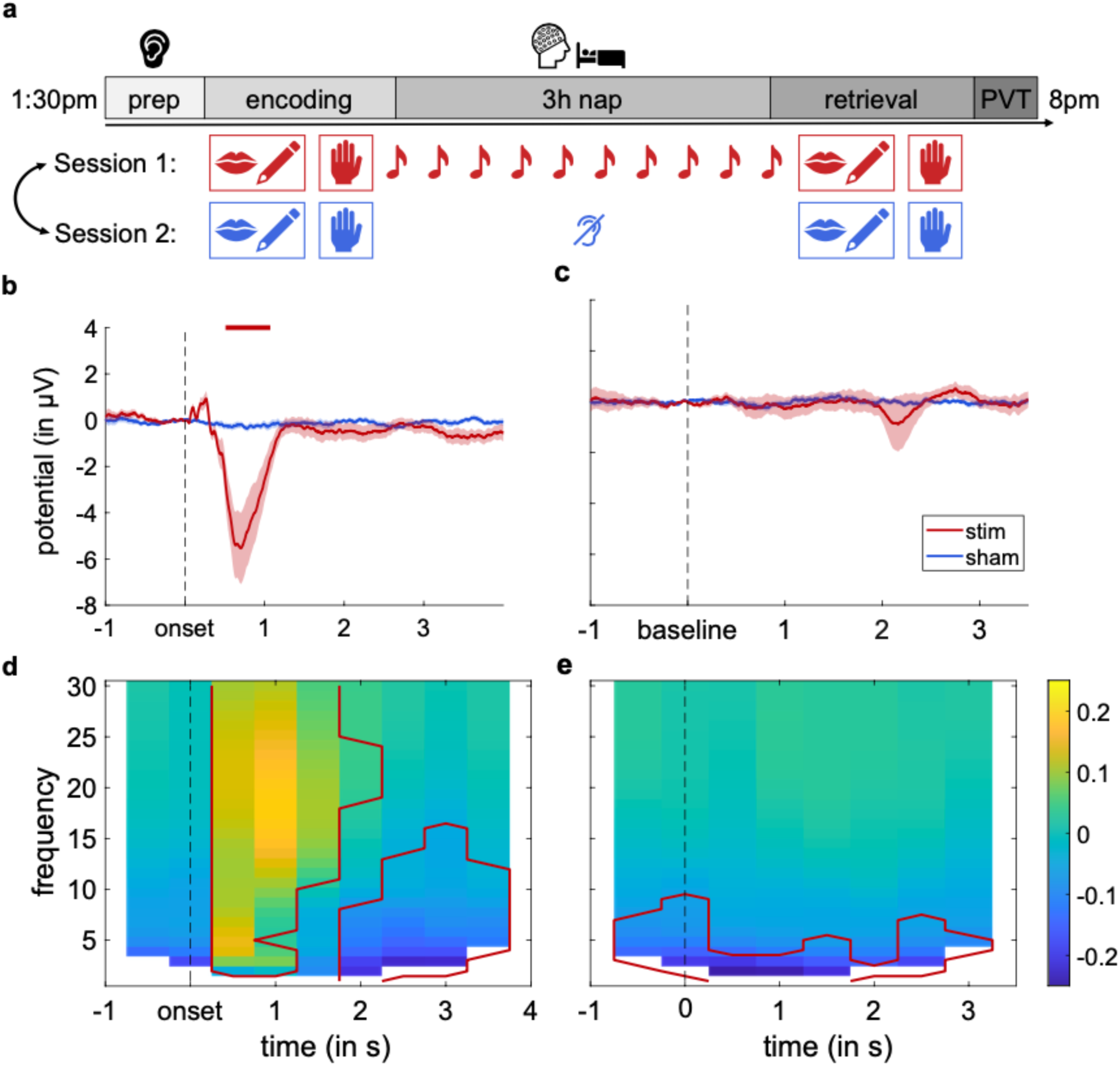
Experimental procedure and auditory stimulation. **a.** After a preparation phase, including hearing tests and EEG setup, participants encoded the declarative (learning-and-memory test, LGT-3) and procedural (sequential fingertapping) memory content. During a 3-h nap opportunity EEG was recorded, and sounds were presented with a random jitter every ∼10 seconds in the stimulation session (red), but not in the sham session (blue). Subsequently, declarative and procedural memory was retrieved. Lastly, a psychomotor-vigilance test (PVT) was performed. The order of stimulation and sham sessions was randomly counterbalanced across participants. **b.** Stimulation triggered a negative event-related potential (ERP). Plot displays average response in frontal channels (Fp1, Fp2, F3, Fz, F4, FC5, FC1, FC2, FC6, C3, Cz, C4, CP5) post sound onset. Time of significant difference shown as red bar. **c.** No ERP differences occurred in segments without stimulation. **d.** Time-frequency (TF) analyses across all channels show a broad band stimulation-triggered increase (yellow hues), followed by a decrease in lower frequencies (dark blue). Red outline denotes significance (p<0.05). **e.** Less power in low frequencies also remained in segments without stimulation.

### Stimulation reduced slow wave sleep, but left sleep duration and general alertness intact

We first compared sleep duration, relative time spent in sleep stages, and vigilance measures to evaluate whether auditory stimulation impacted sleep, whilst retaining comparable alertness, necessary for behavioral contrasts. We observed that the stimulation did not affect sleep duration, measured in epochs scored as asleep (*t*(19) = 1. 91, *p* = .071, *d* = .43; Fig. 2a). However, it significantly reduced the time spent in SWS, both in absolute amount of minutes spent in SWS and relative to the total sleep duration (*t*(19) = 5.37, *p* < .001, *d* = 1.20 and *t*(19) = 5.95, *p* < .001, *d* = 1.33; Fig. 2b). Instead, participants spent relatively more of the sleeping time in the stimulated nap in light non-REM sleep stage N2 (*W* = 89, *p* = .002), even though this effect was not significant for the absolute minutes of sleeping time (*t*(19) = −.78, *p* = .447, *d* = −.17). Importantly, the duration of time spent in N1 and REM stages of sleep did not differ across stimulation conditions (*p* > .620, Supplementary Table S1). Therefore, while auditory stimulation did not prevent subjects from falling asleep or maintaining sleep, it specifically reduced SWS, which was compensated by an increase in lighter N2 sleep. This altered the overall sleep composition while retaining a similar duration. Alertness levels did not mirror the difference in sleep architecture: There was no significant difference in the PVT between the stimulation (*M* = 298.61, *SD* = 24.85, *SE* = 5.56) and sham conditions (*M* = 302.21, *SD* = 25.69, *SE* = 5.74; *t*(19) = −.98, *p* = .338, *d* = .22), nor between the first (*M* = 297.56, *SD* = 25.02, *SE* = 5.59) and second experimental sessions (*M* = 303.27, *SD* = 25.32, *SE* = 5.66; *t*(19) = −1.62, *p* = .121, *d* = −.36).

**Figure 2.**
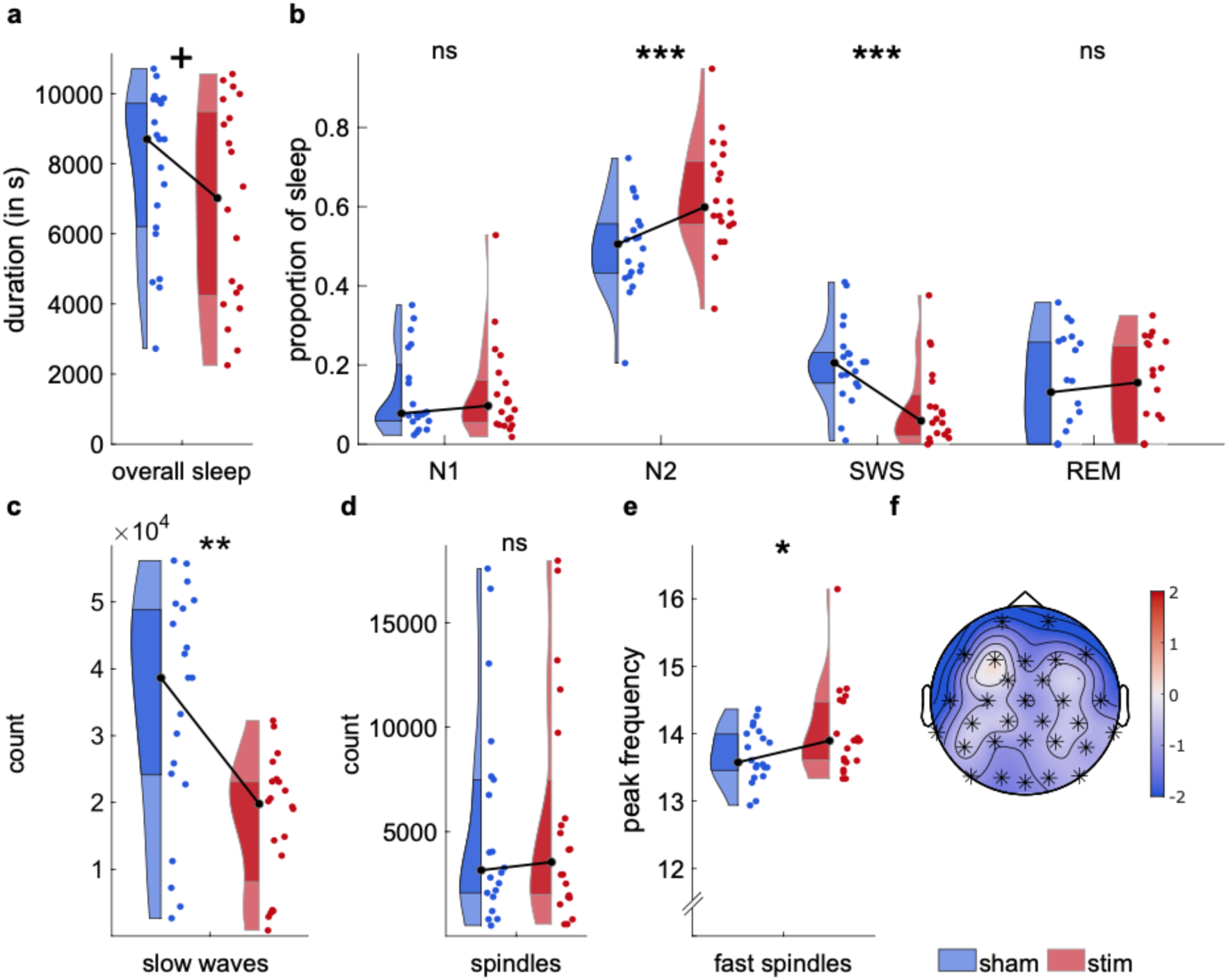
Impact of auditory stimulation on sleep physiology. **a.** Sleep duration (in seconds) in sham (shown in blue) and stim conditions (shown in red). Auditory stimulation did not significantly affect total sleep duration. **b.** Proportion of time spent in each sleep stage, relative to the total duration. Stimulation significantly reduced SWS and increased N2 proportion. **c.** Slow wave count across all channels was reduced for the stimulation condition. **d.** Spindle count did not differ between conditions. **e.** Stimulation was associated with increased peak frequency of fast spindles. **f.** The topography contrast (stim-sham) showed a global decrease in delta power in response to stimulation.

### Stimulation triggered a broad cortical response with lasting low frequency reduction

Since the overall pattern in sleep changed due to stimulation, we also assessed the immediate effects of the auditory stimulus on event-related potentials (ERP) and trigger-bound alterations of the time-frequency trace (TF). Cluster-based permutation testing revealed a significant difference between ERP waves in the stimulation and the sham condition after sound onset, with a frontal cluster (Fp1, Fp2, F3, Fz, F4, FC5, FC1, FC2, FC6, C3, Cz, C4, CP5) showing a negative inflection from .509 to 1.072 seconds post stimulus (*p* = .027, Fig. 1b) and a posterior cluster (TP9, TP10, P7, P8, PO9, O1, Oz, O2, PO10) showing a positive inflection between .375 and .951 seconds post stimulus (*p* = .032). There is no equivalent ERP effect visible between stimulation and sham conditions for segments without stimulation (*p* > .640, Fig. 1c). It should be noted that appearance and features of the triggered ERP waves varied across participants and appeared to be more pronounced during NREM than REM stages (Supplementary Figs. 1 and 2). The stimulation-triggered ERP was matched by the TF analyses, where the stimulation condition showed a significant increase in a broad frequency range (2-30 Hz) and all channels from .5 to 2 seconds post stimulus (*p* < .001, Fig. 1d). Consecutively, 2-3.5 seconds post stimulus, there was a significant decrease (*p* = .019) in lower frequencies (1-16 Hz) across almost all channels (Fp1, Fp2, F7, F3, Fz, F4, F8, FC1, FC2, T7, C3, Cz, C4, T8, TP9, CP5, CP1, CP2, CP6, TP10, P7, P3, Pz, P4, P8, PO9, O1, Oz, O2, PO10). The reduction in lower frequencies (1-9 Hz) observed in the stimulation condition persisted into consecutive time-windows without stimulation (*p* = .035, Fig. 1e).

### Stimulation decreased physiological markers of slow waves but not spindles

Having demonstrated a specific decrease in SWS in the face of stimulation-triggered disruption, we further investigated whether auditory stimulation altered the oscillatory markers involved in memory consolidation, namely slow waves and spindles. Fewer individual slow waves were detected across all channels for the stimulation condition (*M* = 17187.25, *SE* = 2195.45), relative to the sham condition (*M* = 34267.55, SE = 3912.17; *t*(19) = 3.83, *p* = .001, *d* = .86; Fig. 2c). This effect persisted when considering slow wave count during epochs identified as slow-wave sleep only (*t*(19) = 3.92, *p* < .001, *d* = .88). Slow waves also occurred less densely in response to stimulation (*M* = .0044, SE = .0009) as opposed to the sham condition (*M* = .0122, SE = .0014), *t*(19) = 5.74, *p* < .001, *d* = 1.28. In contrast, spindles showed no such detriment of stimulation (*p* > .800, Fig. 2d). However, a further exploratory investigation of potentially affected spindle properties did show a significant increase in peak spindle frequency for fast spindles in the stimulation (*M* = 14.02, *Med* = 13.89, *SE* = .15) in contrast to the sham condition (*M* = 13.68, *Med* = 13.57, *SE* = .09; *W*(19) = 181.5, *p* = .004, *r* = .64; Fig. 2e). No such frequency increase was observed for slow spindles (*W*(19) = 78, *p* = .117, *r* = −.35). There was also no difference across conditions for spindle duration (*p* = .942) or activity (*p* = .486). Likewise, phase-coupling analyses of identified slow waves and spindles showed no differences in mean phases across conditions for frontal or posterior channels (Supplementary Fig. 2). The stimulation cost to slow waves but not spindles was confirmed in frequency analyses: Cluster-based permutation testing on broadband frequency data (0.5-30Hz) revealed a significant negative cluster between stimulation and sham conditions (cluster-level statistic = −1730.1, *p* = .029), amounting to a significant reduction of power almost matching the delta frequency band (.6-5.7 Hz). Analyses for frequencies of interest confirmed significant decreases in delta band (0.5-4 Hz) for the stimulation condition (cluster-level statistic = −41.70, *p* = .008, Fig. 2f), while no condition difference was observed for spindle frequencies (12-16 Hz). Together, stimulation thus consistently affected sleep slow wave activity, reducing slow wave count and density, while leaving markers of spindle activity comparably intact.

### Stimulation disturbed the consolidation of declarative but not procedural memory content

Having confirmed the detrimental effect of auditory stimulation on deep NREM sleep, and specifically slow wave activity, we further assessed whether stimulation impacted declarative and procedural memory consolidation. There was a significant decrease in declarative memory retention, particularly for visuospatial memory, after a nap with auditory stimulation (figural memory score *FS*: *M* = 47.3, *SE* = 2.42), compared to the sham condition (*M* = 53.4, *SE* = 2.27; *t*(19) = 4.35, *p* < .001, *d* = .97; Fig. 3a). However, we found no effect of stimulation on the verbal retention score *VS*, *t*(19) = .36, *p* = .721, *d* = .08, or the overall declarative memory score *DMS* (*t*(19) = 1.34, *p* = .196, *d* = .30; Fig. 3a). Notably, there were no significant declarative memory differences between testing sessions (FS: *t*(19) = .15, *p* = .881, *d* = .04; VS: *t*(19) = −.19, *p* = .848, *d* = −.04; DMS: *t*(19) = −.56, *p* = .581, *d* = −.13; Fig. 3b), confirming the cost to the figural score to be stimulation specific. In contrast to the detrimental effect on declarative memory processing, stimulation did not impact procedural memory performance, assessed in accuracy and speed of reproduced finger tapping sequences (Supplementary Fig. 4).

**Figure 3.**
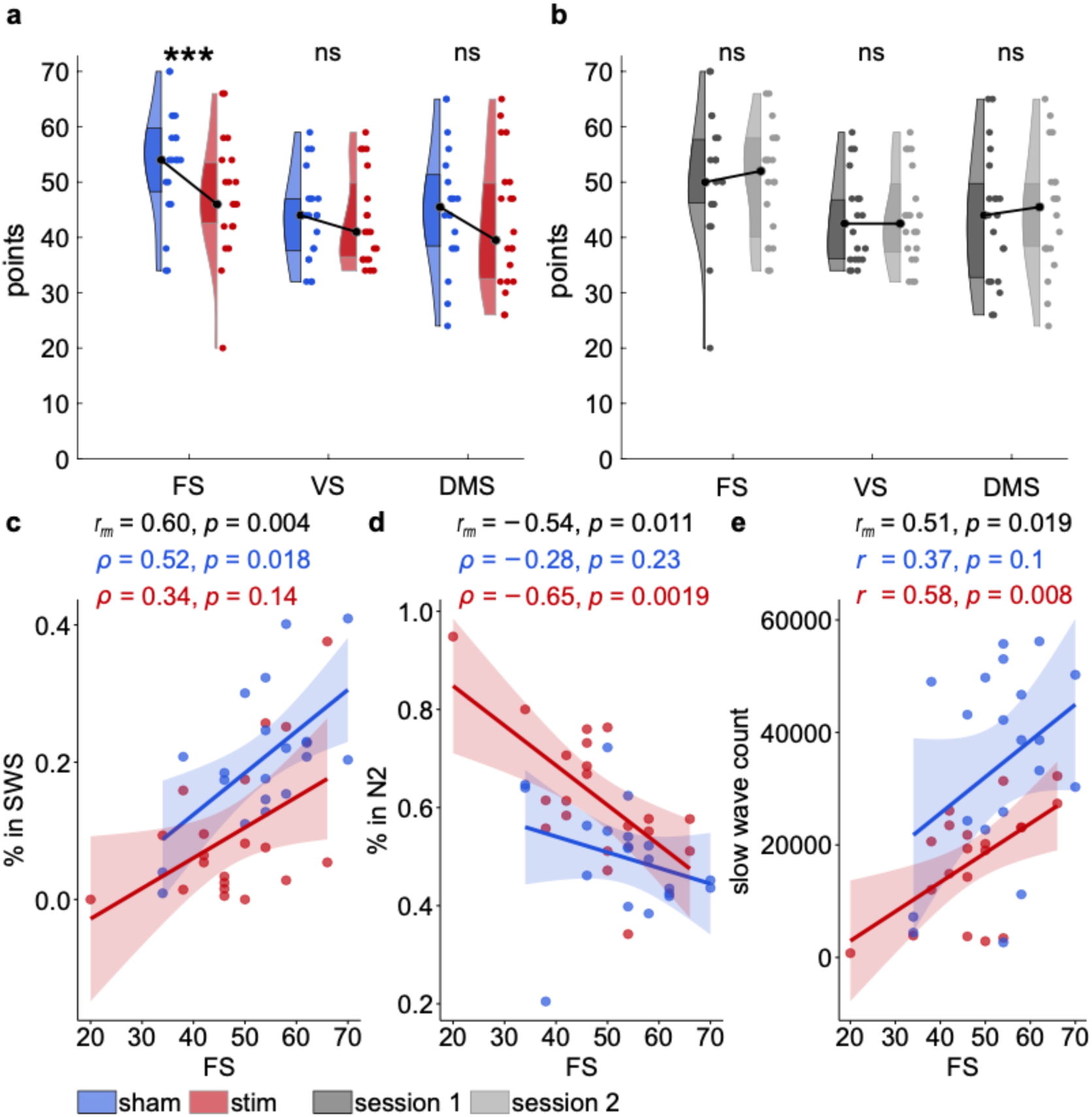
Effects of auditory stimulation on memory performance. **a.** Declarative memory performance, shown as figural, verbal, and overall declarative memory score after the stimulation (red) and non-stimulation nap (blue). Stimulation significantly reduced the figural declarative memory score. **b.** No difference emerged for declarative performance between the first and second experimental sessions irrespective of condition. **c.** Increased proportion of time in SWS was related to improved figural memory performance overall, and specifically in the non-stimulation condition. **d.** Increased proportion of time in N2 was related to decreased figural memory performance overall, and specifically in the stimulation condition. **e.** Higher count of identified slow waves was related to improved figural memory performance overall, and specifically in the stimulation condition. Dots indicate individual data points, line regression of each condition. Black statistics denote repeated measures correlations. Pearson or Spearman correlations were applied depending on normality of analysed data (see Materials and Methods).

### Disruptions in sleep physiology were related to the detriment of declarative memory performance

Considering the impact of auditory stimulation on sleep physiology and declarative memory performance, we tested whether the behavioral and physiological effects were related. Indeed, the differences in sleep architecture in response to stimulation correlated with the figural score, with repeated measures correlations showing a benefit of relative time spent in SWS and a cost of increasing time spent in N2 (*r_rm_* = 0.60, *p* = .004 and *r_rm_* = −0.54, *p* = .011; Fig. 3c, d). Condition-specific correlations reveal that pattern to be driven by a benefit of relative time spent in SWS for the sham condition, *r* = 0.61, *p* = .005 (ρ = 0.52, *p* = .018). In contrast, the relationship between SWS and figural memory performance did not reach significance in the stimulation condition (ρ = 0.34, p = .139). For relative time spent in N2, the detrimental relationship with figural memory was significant for the stimulation condition but did not reach significance in the sham condition (*r* = −0.65, *p* = .002 and *r* = − 0.28, *p* = .226, respectively). Furthermore, greater numbers of detected slow waves positively correlated with higher figural memory scores across conditions (*r_rm_* = 0.51, *p* = .019). Separate analyses showed that this effect was significant for the stimulation but not the sham condition (*r* = 0.58, p = .008 and *r* = 0.37, p = .104, respectively; Fig. 3e).

### Traveling waves as a potential mechanism of propagating information and the limiting effects of stimulation

Given that stimulation had occurred continuously throughout the nap opportunity and significantly impacted sleep architecture and physiology, stimulation may not only have altered the occurrence of slow waves, but also their functional properties. Slow waves are thought to modulate neural excitability while traveling along brain networks and may therefore be a vehicle to propagate information across different neural populations (Massimini et al., 2004; Muller et al., 2018). We identified traveling slow waves by detecting slow waves in each individual electrode and clustering those that occurred spatially adjacent and temporally close into traveling events (Fig. 4a-c, Methods). We found significantly fewer slow wave-clusters indicating traveling slow waves (TSWs) across channels in the stimulation (*M* = 2619.15, *SE* = 319.41) versus the non-stimulation condition (*M* = 5031.25, *SE* = 566.59; *t*(19) = 3.87, *p* = .001, *d* = .87). According to theories of active systems consolidation, slow waves in N2 and SWS are crucial for memory stabilization (Born & Wilhelm, 2012). For the following analyses, we therefore calculated stimulation effects both in reference to all detected slow wave events and additionally restricted to events occurring in these NREM stages only. Indeed, the same pattern of reduced TSWs was observed when only considering only NREM sleep (N2 and SWS; *t*(19) = 3.92, *p* < .001, *d* = .88). Fewer TSWs were expected considering the decrease in both SWS and identified slow waves overall in the stimulation condition. Importantly, the number of identified TSWs correlated positively with behavioral performance in the figural memory task, (*r_rm_* = 0.51, *p* = .018). While this relationship was particularly pronounced in the stimulation condition (*r* = 0.57, *p* = .009), it was not significant in the sham condition (*r* = 0.38, *p* = .098). We observed similar results, when only including waves identified in NREM sleep, i.e., N2 or SWS (overall: *r_rm_* = 0.52, *p* = .017, stim: *r* = 0.55, *p* = .012, sham: *r* = 0.39, *p* = .091). Traveling slow waves thus are of general interest for declarative memory consolidation.

**Figure 4.**
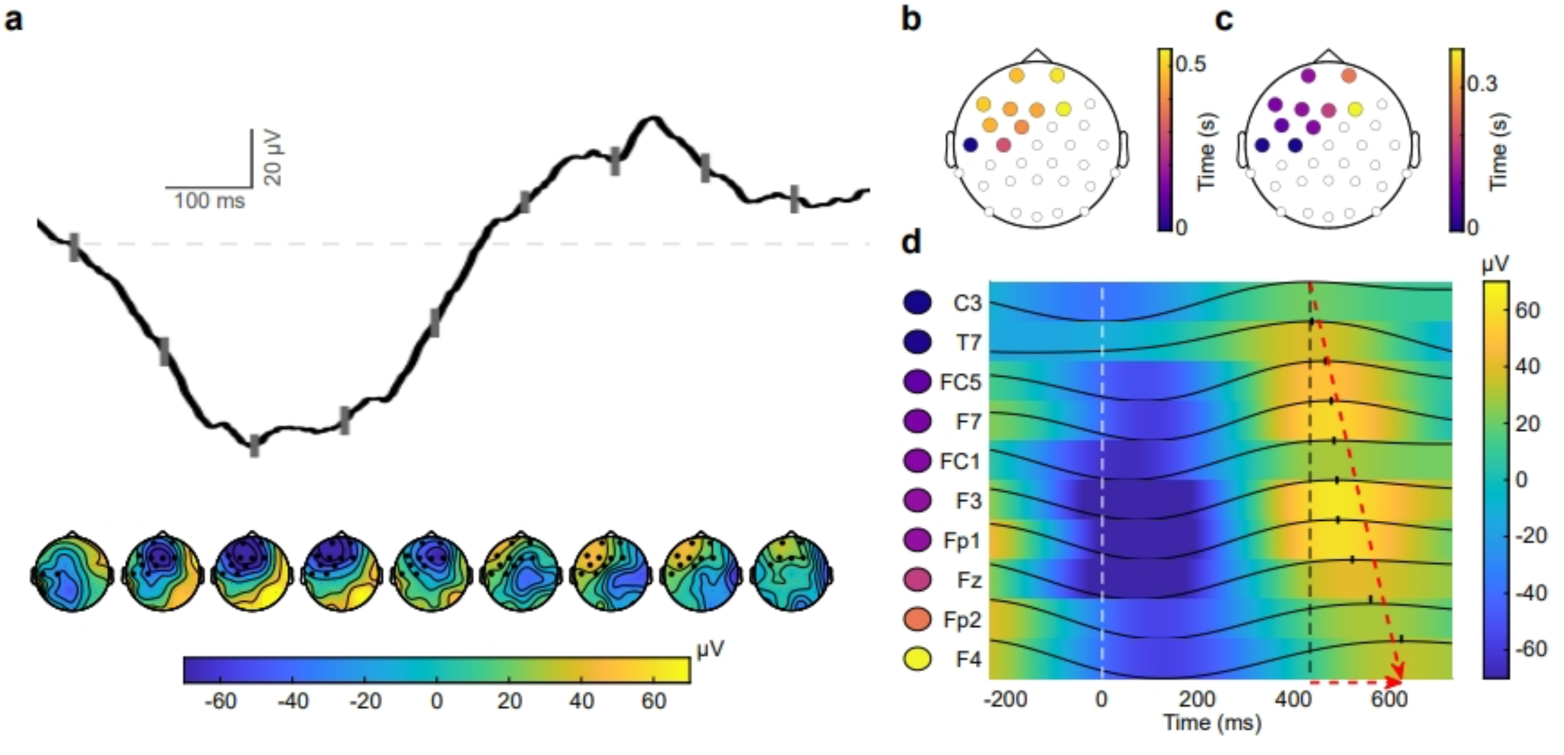
Traveling wave detection. **a.** Exemplary traveling wave with topographical plots illustrating voltage change across time. The waveform was averaged across all involved sensors (highlighted with black circles in the topographies). Gray lines mark timepoints of average voltage shown in the respective topography below. Positive voltage shown in yellow, negative in blue. The spatial trajectory of the wave is illustrated in topographical plots by varying sensor colors for onset time of (**b.**) troughs and (**c.**) peaks of slow waves at each involved sensor. Purple colors denote earlier, yellow colors later onsets. **d.** Averaged waveforms (filtered in the delta-band) for all sensors involved in the exemplary traveling wave, sorted by peak onset times. Markers indicate peak times of each wave and color denotes voltage amplitude (blue negative, yellow positive). Time axis is defined relative to the first trough in the traveling wave. Red arrows indicate the direction and magnitude of peak shift in time across sensors.

### Stimulation altered the nature and trajectory of traveling slow waves

Consequently, we assessed whether stimulation had an influence on the traveling dynamics of the identified slow wave clusters, thus potentially altering the cortical propagation of declarative mnemonic content. We restricted all following analyses of TSWs on non-REM sleep phases N2 and SWS. We contrasted the cortical spread of TSWs across conditions by counting how many TSWs occurred at each sensor pair for the sham and stimulation condition (Fig. 5a). In general, significantly less TSW spread could be observed broadly across the scalp in response to stimulation (Supplementary Fig. 5a). This is unsurprising given the higher TSW count in the sham condition. However, this contrast was significantly enhanced for frontal sensors (Fig. 5a), demonstrating a relatively stronger decrease of TSW spread in frontal regions in response to stimulation. Beyond number and spread, the features of identified TSWs were changed, which extend stimulation effects beyond the known reduction of individual slow waves and time spent in SWS: Peak-to-peak duration of the TSWs was significantly shortened for TSWs in the stimulation (*M* = 97.99 ms, *SE* = 4.48) in contrast to the sham condition (*M* = 111.50 ms, *SE* = 4.68), *t*(19) =2.80, *p* = .011, *d* = .63; Fig. 5b). Across all sleep stages, the overall TSW duration was decreased, and this effect remained at trend level for NREM sleep (*t*(19) = 2.51, *p* = .021, *d* = .56 and *t*(19) = 1.95, *p* = .066, *d* = .44, respectively). Similarly, across all sleep stages identified TSWs also spread to less sensors for the stimulation (*M* = 5.33, *SE* = .21) in contrast to the sham condition (*M* = 5.82, *SE* = .19), and this effect still trended for NREM sleep only (*t*(19) = 2.35, *p* = .030, *d* = .53 and *t*(19) = 2.00, *p* = .060, *d* = .45, respectively; Fig. 5d). The spatial reach of individual TSWs was also hampered by stimulation, showing a shorter maximal distance covered between sensors (*M* = 134.30, *SE* = 2.47) in contrast to sham (*M* = 143.53, *SE* = 2.27; *t*(19) = 3.36, *p* = .003, *d* = .75; Fig. 5f). The reduction of mean travel speed by stimulation remained at trend level (*t*(19) = 2.03, *p* = .057, *d* = .45). The distribution of TSW origins, defined as first to measure the peak of an individual SO in the cluster, was significantly shifted by stimulation, marking less TSW origination from sensor CP5 (*t*(19) = −3.87, *p_BON_* = .033; Fig. 5h). Further considering the spatial distortion of TSWs due to stimulation, we estimated the traveling profiles by percentage of waves engaging different sensor pairs, thus controlling for condition differences in TSW count. Again, we found a significantly broader spatial spread of TSWs across the whole scalp in the sham condition (Supplementary Fig. 5b). However, the main cost of stimulation in terms of TSW spread was particularly focused on frontal sensors (Fig. 5i), confirming our previous results. Overall, stimulation resulted in shorter TSWs with smaller spread to fewer sensors and less spatial expanse, particularly impacting frontal regions.

**Figure 5.**
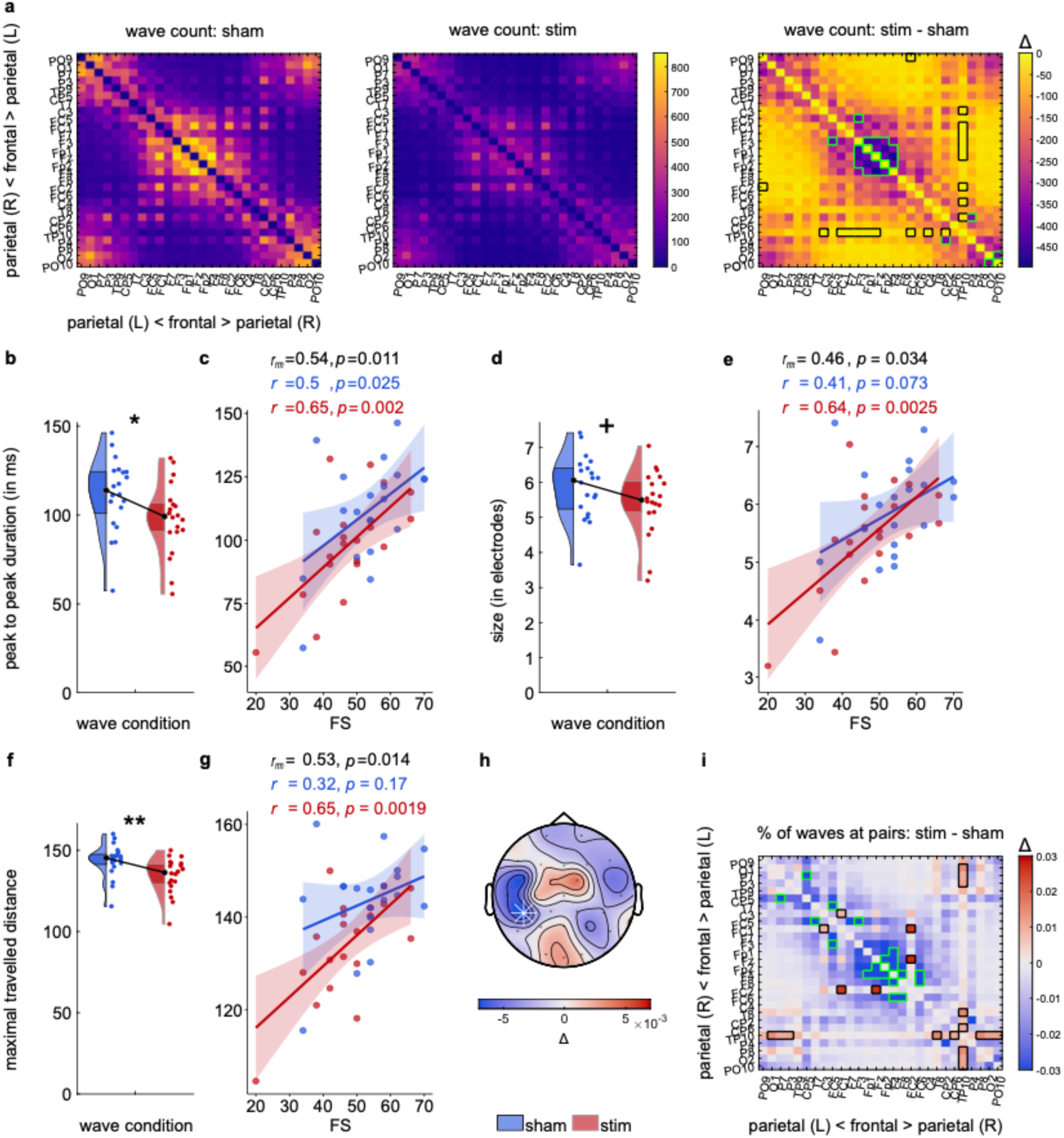
Auditory stimulation disrupts slow wave traveling dynamics. **a.** Count of TSWs involving each sensor pair as measure of cortical spread separately for sham and stimulation session as well as a difference map. Sensors are arranged from posterior parietal on the right via frontal to posterior parietal on the left side of the head. For the individual sessions, yellow color indicates higher count. For the difference, purple color indicates larger differences. Green frames indicate a significantly larger difference, black frames a significantly smaller difference than average. **b.** Duration of wave as maximal temporal peak shift is contrasted for sham (blue) and stimulation (red) conditions. **c.** Peak duration correlated positively with figural score (FS). Dots indicate individual data points, line regression for the condition. **d.** Average number of sensors involved in the TSWs is larger in the sham condition. **e.** The TSW size in sensors positively correlated with the figural score. **f.** Maximal distance covered by TSWs, Euclidean distance between furthest involved sensors, was significantly smaller in response to stimulation. **g.** Larger traveled distance related positively with figural score. **h.** Shift in origin of TSWs due to stimulation. Significant shift away from sensor CP5, indicated by a white star. **i.** Difference in cortical spread of TSWs in each condition (normalized as percent of TSWs involving each sensor pair) indicated broader scalp coverage for TSWs in sham (in blue) but particularly in frontal sensors (green frames). Some posterior sensor pairs showed relatively more involvement for the stimulation condition (in red, significance with black frames). Sensors again arranged from right parietal over frontal to left parietal regions. Black statistics denote repeated measures correlations. Pearson or Spearman correlations were applied depending on normality of analyzed data (see Materials and Methods).

### Alterations of traveling slow waves relate to declarative memory cost

Interestingly, the features of TSWs impacted by stimulation again correlated with behavioral performance in the FS. More precisely, a larger peak-to-peak duration of the TSWs correlated positively with improved figural memory (*r_rm_* = 0.54, *p* = .011), with the relationship holding for both sham and stimulation session individually (Fig. 5c). Furthermore, more involved sensors in the TSWs were related to a higher FS (*r_rm_* = 0.46*, p* = .034), significantly for stim and at trend level for the sham condition (Fig. 5e). Similarly, the distance covered by the TSWs related positively with figural memory (*r_rm_* = 0.53, *p* = .014), even though this relationship was significant only for the stimulation condition (Fig. 5g). Further, the speed with which TSWs traveled, as in duration by maximal cortical spread, was correlated on a trend level with figural memory (*r_rm_* = 0.39, *p* = .080). With respect to condition, the correlation was significant for the stimulation session only (stimulation: ρ = 0.50, *p* = .026, sham: ρ = 0.22, *p* = .350).

## Discussion

Our results unveil a putative relationship between the dynamic features of slow waves and memory consolidation: Beyond the mere presence of slow waves, their traveling trajectories may also contribute to effective declarative memory processing in sleep. We found auditory cues to decrease several markers of SWS quality, leading to lower delta power and fewer emerging slow waves, which was related to impaired declarative memory performance. Furthermore, stimulation decreased the number of traveling slow waves, with the remaining events being restricted in duration, size, and spatial expanse. Importantly, these features of traveling slow waves were related to declarative memory consolidation, indicating that sounds presented during sleep prevent successful memory processing not only by suppressing slow wave activity, but also by hindering the natural propagation of occurring slow waves.

The observation that sounds during sleep came at a cost to sleep depth rather than overall duration replicates findings in the literature (Fehér et al., 2023; Muzet, 2007; Schönauer et al., 2014). Specifically, auditory stimulation appeared to interfere with entering and maintaining slow wave sleep, instead resulting in a higher proportion of N2 sleep. This physiological alteration could be triggered by different, complementary mechanisms: The stimulation-dependent decrease in slow wave density might have prevented epochs from reaching the necessary scoring criteria of 20% slow waves. Also, auditory stimulation could trigger an arousal response resulting in a higher likelihood of moving to lighter sleep with fewer slow waves (see below). Importantly, the relative increase in N2 could account for the observed lack of stimulation-induced spindle reduction, given that spindles occur in both SWS and N2. Taken together, our findings that auditory stimulation decreased SWS, disrupted slow waves, and impaired declarative memory consolidation support theories of active systems consolidation during sleep (Born & Wilhelm, 2012; Klinzing et al., 2019; Staresina et al., 2015), which stress the importance of a coordinated interplay between slow waves, spindles, and ripples for declarative memory processing. This importance of slow waves for declarative memory consolidation is supported by studies emphasizing mnemonic gain or loss depending on slow wave increase (Marshall et al., 2006; Ngo et al., 2013; Westerberg et al., 2015 but see Cordi & Rasch, 2021) or decrease (Backhaus et al., 2007; Mander et al., 2015).

Our results further suggest that memory consolidation depends not only on the occurrence of slow waves, but also their dynamic trajectory and profile across the scalp: Stimulation stunted the expanse of the traveling slow waves in terms of involved sensors, duration, and physical reach across the scalp. These altered features in turn correlated with the impaired memory performance, indicating that the dynamic features of traveling slow waves have a role in declarative memory consolidation. In general, traveling waves observed across different frequency bands and spatial regions have been associated with various processes involved in computation and information propagation (Muller et al., 2018): For example, hippocampal theta oscillations do not just represent behavioral information but also exist as traveling waves (Lubenov & Siapas, 2009; Zhang & Jacobs, 2015), with their propagation dynamics relating to behavioral task performance (Zhang et al., 2018). Spindles, which have been shown to travel globally across brain regions along preferred spatiotemporal paths (Muller et al., 2016), demonstrate highly organized neuronal co-firing depending on their propagation pattern, arguably preparing plasticity in local cortical regions (Dickey et al., 2021). Slow waves can also travel in complex cortical patterns (Hangya et al., 2011; Massimini et al., 2004) and have been associated with information transfer across brain hemispheres (Avvenuti et al., 2020; Bernardi et al., 2021). Indeed, changes in traveling profiles of slow waves during development from childhood to adulthood related to brain-wide connectivity (Castelnovo et al., 2023), whilst in the aging brain, tau pathology observed in diseases affecting long-term memory stunted the traveling profiles of slow waves (Sharon et al., 2024). Interestingly, during memory tasks, different traveling patterns of waves have been associated with particular cognitive processes or mnemonic content (Das et al., 2024). The functional role of traveling waves in cross-regional connectivity and (mnemonic) information transfer aligns with our interpretation that the dynamic traveling profile of sleep slow waves facilitates the propagation of information necessary for successful memory consolidation during sleep. We find that experimentally disrupting SWS, including the occurrence and traveling properties of slow waves, impairs memory performance. Given that auditory stimulation affected both the overall count as well as the characteristics of individual slow waves, we can only provide correlational evidence that both number and expression of traveling slow waves are involved in declarative memory consolidation. In the future, we may be able to disentangle the specific contribution of the propagation of mnemonic information and slow wave generation, more generally. Potentially, time-locked stimulation (Fehér et al., 2023; Ngo et al., 2013) could be implemented to disrupt slow wave activity with a specific focus on dynamic features while retaining sleep depth.

Although we observed the different features of traveling slow waves were related to declarative memory consolidation, we did not observe a detrimental effect of auditory stimulation on procedural memory consolidation. Sleep-dependent processing of procedural memories has frequently been linked to spindle activity (Kumral et al., 2023; Nishida & Walker, 2007; Wamsley et al., 2012). Therefore, the specific decrease in SWS should not affect the processing of procedural content, especially if compensated by an increase in N2 (Walker et al., 2002). The dissociation between declarative cost and procedural stability corroborates findings in the literature speaking to a dissociation between the sleep-related consolidation processes of declarative and procedural memories: Plihal and Born (1997) observed a larger reliance of declarative processing (paired-associate lists) on SWS-heavy early night sleep, whilst procedural tasks (mirror tracing) benefited more from later night sleep, predominant in N2 and REM phases. Increasing SWS using a GABA agonist can actually deteriorate procedural memory consolidation (Feld et al., 2013; Walker et al., 2002). Overall, the shift from SWS to lighter sleep stages caused by the auditory stimulation appears to affect predominantly declarative memory processing, thereby sparing procedural memory processing.

One factor contributing to reduced deep sleep in the stimulation condition could be arousal following the auditory cue. Salient sensory input during sleep can evoke k-complexes, which may represent either an arousal or gating mechanism on whether the information needs to be processed and an awakening should follow (Halász, 1993, 2016). In this study, we observed a broad cortical response time-locked to the stimulation, indicating the triggering of k-complexes. Sound-induced arousal mediated by k-complexes might lead to a transition from slow-wave sleep to lighter sleep, such as N2 (Fehér et al., 2021; Lechat et al., 2021). Interestingly, some authors have also made a distinction between type I (k complex-like) and type II (homeostatically regulated) slow waves (Bernardi et al., 2018): The auditory stimulation in the current study may have triggered type I slow waves and decreased the occurrence of type II slow waves. Our analyses of the traveling wave forms support this idea: On average, the detected traveling waves exhibited a shorter duration for the stimulation condition, similar to the difference observed between type I and type II waves. Type I waves are also associated with an increase in high-frequency activity, which could be observed in the stimulation condition. Subsequently, power in the lower frequency bands, encompassing delta, theta and spindle band were suppressed, which is coherent with the previously reported type I wave-bound suppression of the spindle frequency band (e.g., see Bernardi et al., 2018). Importantly, spindle analyses provide further support for the arousal hypothesis, by showing that stimulation caused an increase in peak spindle frequency for fast spindles. Previous research has linked the speed of peak spindle frequencies to sleep depth, indicating faster spindles during nap periods compared to night sleep (Bódizs et al., 2022) and during lighter sleep versus deep sleep (Andrillon et al., 2011; Siclari et al., 2014). An increase in peak frequency of fast spindles for auditory stimulation could thus be an indicator of lighter sleep or arousal. The stimulation-related increase in peak fast spindle frequency is also consistent with the observation that externally elicited spindles are faster than spontaneously occurring ones (Klinzing et al., 2021). In accordance with the present study, Kokkinos and Kostopoulos (2011) also found that while k complexes interrupted concurrent spindles, subsequent spindles increase in frequency by about 1 Hertz. Interestingly, Cairney and colleagues (2018) report an increase in fast spindles in response to auditory TMR and Ngo and colleagues (2013) in response to auditory stimulation in-phase with slow waves. Thus, auditory stimulation during sleep may cause arousal responses even if they are timed to the slow wave phase or act as memory reactivation cues.

The arousal(-like) response to auditory stimulation during sleep contrasts with the success of TMR and closed-loop stimulation of slow waves in promoting memory consolidation and thus raises both functional and methodological questions. Firstly, given the ostensible success of auditory TMR to boost memory retention (Oudiette & Paller, 2013; Rudoy et al., 2009; meta-analysis: Hu et al., 2020), it could be possible that content-loaded stimuli are processed differently to random noises, or that the gain derived from triggering a reactivation event outweighs the arousal costs, leading to a net benefit. The increased response strength in the spindle frequency band evoked by memory-associated cues compared to neutral cues supports the idea that post-cue processing depends on stimulus content (Cairney et al., 2018). Further studies are needed that explicitly manipulate the balance of costs and benefits for presenting content-loaded auditory stimuli during sleep. Secondly, the specific impact of the auditory stimulation is likely bound by its temporal relation to ongoing slow wave activity. Closed-loop algorithms stimulating during the up-phase of the slow wave have been shown to enhance slow wave activity (Besedovsky et al., 2017; Esfahani et al., 2023; Fehér et al., 2021; H.-V. V. Ngo et al., 2013). Importantly, Ngo and colleagues observed that the traveling profiles of spontaneously occurring and induced slow waves, i.e., stimulated when the next predicted slow wave up-phase should occur, were comparable across a range of measures. Combined with our finding of randomly timed stimulation disturbing both slow wave activity and their usual traveling profiles, it appears that the precision of the algorithm timing may underlie this advantage of closed-loop stimulation. Indeed, using detection algorithms to stimulate slow waves out of phase can successfully disrupt slow wave activity and thereby suppress slow-wave sleep in humans (Fehér et al., 2023). Interestingly, this closed-loop suppression produced a comparable reduction in SWS to the random auditory stimulation performed in this study. Therefore, at least for SWS suppression, random auditory stimulation might be a suitable, simplified alternative to phase-locked stimulation. Developing a simple and reliable method to disrupt slow-wave sleep has clinical and experimental value, such as exploring the role of SWS in emotional memory processing or modeling sleep fragmentation in different neurological disorders. The negative effects of random noise on sleep and cognition are also relevant to society, as sleep duration alone is not a valuable predictor for sleep health: also sleep depth and the properties of sleep oscillatory activity need to be considered.

In conclusion, our results critically extend existing literature on auditory stimulation in sleep by showing that random stimulation disturbs specifically SWS, slow wave occurrence, and their traveling profiles, at the cost of declarative memory consolidation. Whilst supporting current theories of sleep-dependent memory consolidation, our findings highlight the need to consider dynamic features of sleep physiology beyond average markers like sleep duration. Furthermore, the observation that stimulation untied to learning content or not timed to ongoing oscillatory activity decreases memory performance has methodological implications for the use of targeted memory reaction and time-locked stimulation paradigms.

## Acknowledgements

We thank Susanne Möhrle for help with data acquisition and Angelina Eisele for her contribution to data visualization. This work was supported by Deutsche Forschungsgemeinschaft (DFG, German Research Foundation; grant: SCHO1820/2-1, grant: GA730/3-1, grant RO6828/1-1).

## Author contributions

S.G. and M.S. contributed to the design and implementation of the research. N.M.R. completed data preparation, statistical analyses and writing of the manuscript. D.K. completed the phase coupling analysis and provided feedback on the manuscript. M.S. further contributed to writing of the manuscript.

## Competing interest statement

The authors have no conflicts of interest to declare.

## Materials and Methods

### Participants

Twenty-five healthy male and native German-speaking adults were recruited for this study. All participants were non-smokers, right-handed (as assessed by the Edinburgh Handedness Scale, Oldfield, 1971), and had no diagnosed sleep-or memory-disorders. None consumed drugs or were prescribed medication that could affect their central nervous system. The participants had a regular sleep rhythm of 7-9 hours (measured by the Munich Chronotype Questionnaire, MCTQ, Roenneberg et al., 2007) and did not work in shifts or experienced jet lag in the last six weeks before the study took part. Further task-specific exclusion criteria were applied: Piano players and Turkish speakers were excluded to ensure measure efficacy in the finger tapping and LGT-3 tasks, respectively. Distinct and superior finger dexterity in piano players (e.g., Furuya & Altenmüller, 2013) would bias performance, given that finger tapping is similar to piano playing. The LGT-3 includes learning vocabulary in a foreign language (Turkish), which would become redundant if the language was already known.

Five participants had to be excluded due to withdrawing from the study prior to the second session (n=2), equipment failure in recording electrophysiology continuously (n=2) or due to a failure to fall asleep in an experimental session, defined as at least 20 minutes in sleep stage N2 or deeper (n=1). All analyses were performed on the data of the remaining set of twenty participants (age: *M* = 24.05, *SD* = 3.91, range: 18-31).

### Procedure

The study was carried out on two experimental visits, separated by at least six days (*M* = 17.95, *SD* = 12.17), each including declarative and procedural memory tasks as well as a three-hour sleep period (nap). The task versions and the order of the stimulation conditions during the nap were randomized and counterbalanced across participants (Fig. 1a). Participants were recompensed for their participation at a rate of 8 Euros per hour.

In preparation for the study, participants completed the Munich Chronotype Questionnaire (Roenneberg et al., 2007) to evaluate regular sleeping behavior. In order to increase the likelihood of sleep during the nap opportunity, participants were asked to go to bed regularly the three preceding nights but rise an hour earlier than usual. Additionally, participants were asked to refrain from consuming caffeine or medication on the experimental days as well as alcohol on the days of and preceding the experiment. A hearing test was administered to assess participants’ hearing threshold prior to the experiment. The encoding session was equivalent for both experimental visits, consisting of one version of the LGT-3, followed by one type of sequence learnt in the finger tapping task.

Participants then had a three-hour nap opportunity, during which EEG was recorded. In the stimulation condition, a train of six clicks (10 Hz stimulation train, lasting 600 ms) was presented every 9.5 seconds plus jitter (*M* = 9.5313, *SD* = .0013) via in-ear headphones. The volume of the stimulation was adjusted by the participants with the instruction to keep it as high as possible and as low as necessary, since they should be able to fall asleep. Experimenters ensured that the chosen level was above their individual hearing threshold and above the minimum audible level (hearing threshold *median* = 29.3; *SD* = 11.6; stimulation intensity *M* = 72.12, *SD* = 7.31 [calculated decibel level directly at sound source, lower intensity at ear]). No sounds were presented during the sham condition. After the nap opportunity, participants completed the retrieval phase, consisting of the recall of the declarative information learnt in the LGT-3 set and the performance of the finger tapping task. At the end of each experimental day, a psychomotor vigilance task was performed.

### Task material

#### Hearing test

The hearing test was completed in the first session only. Via Matlab, a continuous sinus tone with a frequency of 4000 Hz was presented, which steadily increased or decreased in volume. Participants had to press the “up” or “down” key respectively, if they could hear it for the first time (getting louder) or for the last time (getting quieter). Increase and decrease trials were sampled in random order. Hearing threshold was assessed as the median individual measured output source values (*median* = 29.3, *SD* = 11.6 [calculated decibel level directly at sound source, lower intensity at ear]). Hearing threshold values were used to confirm that participants were able to perceive the acoustic stimulation during sleep.

### Learning-and-Memory-task (Lern-und-Gedächtnistest, LGT-3)

The LGT-3 is a German test battery of different paper-and-pencil declarative memory tasks (Bäumler, 1974), assessing a total declarative memory score (DMS), as well as the subcomponents figural memory (figural score, FS) and verbal memory (verbal score, VS). The individual tasks consist of two figural tasks comprising the figural score (spatial navigation on a map, identifying fictional brand logos), three verbal tasks combined for the verbal score (German-Turkish word-pairs, memorizing phone numbers, recalling content from building plans such as terminology or numbers), and an object memory task. The different scores are calculated as the weighted sum of relevant task components. In this study, the two equivalently difficult task sets were randomly allocated across stimulation and sham condition. The test procedure was adapted to accommodate the three-hour nap opportunity between encoding and retrieval phase. Prior studies suggested that this delay did not hamper the measurement efficacy, and that task performance benefits from sleep in contrast to a wake condition (Schönauer et al., 2014).

#### Finger tapping

Procedural learning was assessed via a finger tapping task, which is sleep-dependent (Walker et al., 2003, 2005). The motor task consisted of a sequence of five numbers that participants had to tap on a keyboard in the right order. There were two equivalently difficult versions of the sequence (FT1: “4-1-3-2-4”, FT2: “4-2-3-1-4”) (Walker et al., 2003), and each study day was randomly assigned one version. In a fixed task window (“block”) of 30 seconds, participants were tasked to type the chosen sequence correctly and as often as possible, using their non-dominant hand (i.e., left in this study). For this, participants were instructed to focus on both speed and accuracy. Outcome measures per block are mean time until button presses (reaction times, “RT”, in ms), the total number of typed sequences (“count”), as well as the number of correctly typed sequences (“accuracy”). The encoding phase consisted of 12 blocks, separated by 20-second-long recovery breaks. The last three blocks of the encoding phase are used for statistical assessment, to match the three blocks implemented as the retrieval phase.

The finger tapping task was computer based with a generic keyboard and presented via Matlab. The participants were in a darkened room and focussed on the screen displaying the sequence in white script on black background. After typing a number, a star symbol (*) appeared, indicating the current position within the sequence to the participants. Auditory feedback was provided via a speaker for wrong key presses. The participants also received an auditory alert paired with a visual prompt to indicate the start of the next block after each recovery break.

#### Psychomotor vigilance task (PVT)

This simple visual reaction time test (Dinges & Powell, 1985; Drummond et al., 2005) was implemented to monitor alertness and vigilance for each experimental session. Four red numbers were presented on a darkened computer screen and started counting upwards in milliseconds at irregular time intervals. Participants were instructed to press the spacebar to stop the counting as quickly as possible with their dominant (right) hand as soon as they saw the numbers changing. The resulting number on the screen represented their reaction time in milliseconds. Button presses <200ms counted as wrong (impulsive) presses, and presses >500ms as lapses. The wrong types of presses were analyzed separately from the rest.

#### Electrophysiology recordings and preprocessing

Electroencephalography (EEG) was measured with a standard setup (10-20-System, Kasper, 1985) and amplifier by Brain Products, using Ag-AgCl-electrodes in the ActiCap 128 electrode system. The EEG recording was measured with 32 channels and a sampling rate of 1000Hz during the sleep period. Separate derivations for EEG, electrooculogram (EOG) and electromyogram (EMG) derivations of the complete data were created and scored in 30-second windows according to Rechtschaffen and Kales (1968) by two expert scorers. Discrepant scoring was resolved by a third expert scorer. Data preprocessing was completed using Fieldtrip (Version: 20230418, Oostenveld et al., 2011), initially demeaning, detrending and filtering the data (low-pass: 140 Hz, high-pass: 0.1 Hz, notch: 50, 100, 150 Hz) each channel. Trials were created locked to stimulation times or equivalent time stamps for the non-stimulation condition and used for artifact rejection: After a manual rejection of channels and trials based on variance, EOG, jump, and motion artifacts were detected based on adjusted thresholds and cut from the data. Missing channels were interpolated using the spline method before the cleaned data were then re-referenced to the average reference.

##### Frequency and time-frequency analyses

Frequency and time-frequency analyses of overall and stimulation-locked activity were performed using Fieldtrip. To achieve higher frequency accuracy especially for lower frequency, both analyses were completed separately. Frequency was estimated by using the entire data implementing a Hanning taper for the frequency band of 0.5 to 30 Hz. Statistical contrasts across conditions were calculated using cluster-based permutation testing across all frequencies, and specifically for delta and spindle frequencies (averaged across frequency band). Time-frequency decomposition of the stimulation-locked data (−1s to 4s, relative to stimulation) was achieved using a Fourier analysis based on sliding time windows in steps of 50ms, also implementing a Hanning taper. The window length was set to three cycles of the given frequency (1 to 30 Hz), moving in 1 Hz steps. Statistical analysis was also performed using cluster-based statistics, either across all frequencies and time-points post stimulation, or for frequencies of interest (averaging across spindle or delta band). All Fieldtrip cluster-based permutation tests (also for the following analyses) are inherently corrected for multiple comparisons.

##### Event-related potential analyses

Using Fieldtrip, the trials time-locked to stimulation times (−1s to 4s, relative to stimulation) were detrended and a baseline was applied for 100ms before stimulation onset. Event-related potentials were then calculated, averaged across trials, and statistically evaluated with cluster-based permutation testing.

##### Spindle peak-frequency

The power spectral density (PSD) was estimated using the MATLAB pwelch function, restricted to frequency range of 0.1 to 45 Hz. Fast Fourier Transform was computed on the signal divided into overlapping segments (by 95%) with window length of twice the sample length. The average, log-transformed PSD was then used to find peaks in the fast (13-17Hz) and slow (11-13Hz) spindle range. In accordance with literature (Andrillon et al., 2011), fast spindles were defined as beyond 13 Hz and have been associated with posterior topography (Cox et al., 2017).

##### Spindle and slow wave detection

Cleaned trials were concatenated to create a pseudo-continuous artifact-free file, before being bandpass filtered to either spindle (11-17 Hz) or slow wave (0.5-4 Hz) frequencies. Detection algorithms were processed for each channel. The spindle detection algorithm creates an envelope around the spindle-filtered data, and identifies peaks in the envelope (minimum distance: 25ms). The amplitude criterion was defined as exceeding the 95th percentile relative to REM peaks if present, otherwise on N1 peaks. Spindles were identified if exceeding the amplitude criterion for 0.5 seconds, and duplicates were deleted in favor of the maximal option. Parts of spindle events crossing non-continuous trial boundaries were excluded, unless the spindle would then not fulfill the duration criterion, in which case the whole spindle was excluded. The slow-wave detection defined peaks and troughs as well as inflection points. Identified slow waves were excluded if they crossed non-continuous trial boundaries or were below the amplitude criterion, defined as the 99th percentile of N1 sleep peaks. Slow wave density was defined as the count of slow waves in SWS, divided by the number of SWS epochs contributing to the count.

##### Slow wave clustering

Identified slow waves across channels were sorted by peak time. Using the fieldtrip neighborhood structure method triangulation, we identified relevant neighboring sensors for each EEG sensor, and grouped individual slow waves in clusters if peaks fell both within neighboring structures and within 200ms of each other (Kurth et al., 2017; Schoch et al., 2018). Only clusters containing at least two sensors were included. Using surface EEG, the traveling trajectory of the slow wave can be observed by plotting scalp topographies at various points of the average waveform, illustrating the progressive voltage shift across sensors and time (Fig. 4a). The spatial progression of the wave across sensor space is illustrated more clearly by coloring the wave troughs (Fig. 4b) and peaks (Fig. 4c) by onset time. The temporal shift of negative and positive peaks becomes evident when contrasting the waveforms and voltages for each sensor, demonstrating both the cumulative voltage change in sensors as well as the temporal shift of peak to peak across individual waveforms (Fig. 4d).

##### Wave parameters and statistical analysis

Overall wave duration measured the difference between the first and last instance of any slow wave within the cluster, using the downward inflection points before and after trough and peak, respectively. Peak duration represents the maximum temporal shift of peaks within the clusters, thus the difference between first and last maximum of individual slow waves. The size of the cluster was defined by how many slow waves were grouped within the cluster. The maximum distance covered was calculated by the Euclidean distance between the two sensors in the cluster furthest apart. Speed was estimated by the quotient of maximal distance and total wave duration. Statistical analyses were performed in R, using t-tests or Wilcoxon Signed-Rank tests. The origin of the traveling wave was defined as the sensor with the first peak of the individual slow waves in the cluster.

The distribution of origins across sensors was then normalized by total count of clusters within the respective condition. We used a mass univariate approach of statistical analysis, comparing normalized frequency of origin for each sensor, and using Bonferroni adjustments to correct for multiple comparisons. The spread of the wave was estimated in two ways: One measure was a simple count of waves present for each electrode pair. The other measure normalized the count distribution by the total count of clusters in the respective condition. For statistical analysis, a vector was created containing the across-subjects average difference of (relative) counts between conditions for each unique sensor pair (i.e., below the diagonal). This vector contained the real measured condition difference and was contrasted with two different types of permutation distributions, the first shuffling condition labels and the second shuffling all obtained stim-sham difference values: In the first assessment, random difference vectors are created across 1000 permutations by flipping the condition labels randomly for each subject and taking the mean difference between the randomly flipped conditions across subjects. This approach highlights whether and where there are differences across conditions. In the second analysis, the real difference vector was shuffled 1000 times across locations to create a random value distribution for each sensor pair. This approach highlights where existing condition differences were particularly prominent. For both options, each actual value was deemed significant at an alpha of .05, if it was within the 2.5% extreme tail ends of the randomly permuted distribution for that sensor pair. Significance is portrayed mirrored for both representations of each sensor pair.

##### Wave parameters and statistical analysis

Unless otherwise specified, statistical analyses were performed in R, using t-tests or Wilcoxon Signed-Rank tests as appropriate. Relationships between sleep parameters and behavioral measures were assessed using Pearson or Spearman correlations, depending on the normality of the underlying data distributions. Repeated measures correlations (‘rmcorr’, Bakdash & Marusich, 2017; signified as *r_rm_*) were used to assess relationships of variables across conditions, thereby containing multiple measures per participant.

## Supplementary material

### ERP differences across NREM and REM

The discussed ERP effects and individual subject data indicated that auditory stimulation triggered k complexes, even though not all subjects showed them to the same extent or at all: Eleven subjects showed immediate evoked responses, seven subjects showed no evoked potentials, and two subjects showed a delayed reaction (examples shown in Supplementary Fig. 1a, c). Importantly, the amplitude of the k complex-like potentials appeared larger in NREM sleep stages (N2 and SWS) in contrast to REM (Supplementary Fig. 1a and c). It is thus unsurprising that the stimulation only resulted in significant ERPs for NREM sleep (*p* = .012, Supplementary Fig. 1b) and not for REM sleep (*p* = .239, Supplementary Fig. 1d). It should be noted that only eleven subjects had REM sleep in both nap sessions and could thus be statistically compared.

### Evoked-response related analyses

Given the differential evoked responses to the stimulation, we further analyzed the stimulation-dependent effects on sleep architecture and declarative memory based on whether the participant showed some evoked response (n=13) or not (n=7). Both evoked and non-evoked groups show no significant difference in overall sleep duration (*t(12)* = 1.89, *p* = .083, *d* = .52 and *t(6)* = .48, *p* = .180, *d* = .48; Supplementary Fig. 2a, c). Stimulation presented with more relative time spent in N2 (evoked: *W* = 85, *p* = .003, non-evoked: *t(6)* = −3.45, *p* = .014, *d* = −1.31) and less in SWS in both (evoked: *t(12)* = 4.91, *p* < .001, *d* = 1.36, non-evoked: *t(6)* = 3.30, *p* = .016, *d* = 1.25; Supplementary Fig. 2b, d). Again no differences were observed in N1 (evoked: *W* = 56, *p* = .497, non-evoked: *t(6)* = 1.16, *p* = .290, *d* = .44) or REM (evoked: *t(12)* = .50, *p* = .625, *d* = .14, non-evoked: *W* = 9, *p* = .787). Similarly, both subgroups showed a significant reduction in figural memory in the stimulated condition (evoked: *t(12)* = −3.51, *p* = .004, *d* = .97, non-evoked: *t(6)* = −2.46, *p* = .049, *d* = .93; Supplementary Fig. 2e, f). No effects were observable for verbal memory (evoked: *t(12)* = −.38, *p* = .711, *d* = .11, non-evoked: *t(6)* = −.05, *p* = .959, *d* = .02) or the overall declarative memory score (evoked: *t(12)* = −1.17, *p* = .264, *d* = .32, non-evoked: *t(6)* = −.66, *p* = .534, *d* = .25). In summary, whilst the effects on sleep architecture and declarative memory appear to be more pronounced in participants reacting to stimulation with an evoked response, stimulation also had a similar effect in subjects without such response.

### Phase coupling analyses of slow waves and spindles

Phase information for each individual slow wave was derived from the detected peak, trough, and inflection points (see Methods). Accordingly, we detected spindles occurring within each individual SO and aligned them with their respective phases. The phase information for each SO-spindle onset pair was initially averaged across the regions of interest (ROIs; Frontal: F3, Fz, F4; Parietal: P3, Pz, P4) for each participant. It should be noted that, due to the reduced number of identified slow waves in the stimulated condition, there were also fewer slow wave-spindle pairings identified across the ROIs in the stimulated (frontal: *M* =176.30, SE =42.56, parietal: *M* = 82.40, SE = 19.86) than in the sham condition (frontal: *M* = 198.60, SE = 45.18, parietal *M* = 86.60, SE = 25.50).

We used circular statistics, specifically the Hotelling paired sample test for equal angular means, to further compare conditions. Though numerically smaller, the mean phase of SO-spindle pairs in the stimulated (*M* = 188.21_°_, *SE* =15.77) was not significantly different from the sham condition (*M* = 175.17_°_, *SE* =15.66) in parietal regions (*F*(19) = 0.01, *p* = .99; Supplementary Fig. 3a). For frontal regions, the difference between stimulated (*M* = 203.34_°_, *SE* =8.00) and sham condition (*M* = 215.74_°_, *SE* =8.38) was more pronounced, but did not reach significance (*F*(19) = 1.69, *p* = 0.21; Supplementary Fig. 3b). Our results thus indicate spindle onset to occur earlier in the slow wave phase in the stimulated condition.

### Procedural memory analyses

Stimulation did not significantly impact procedural memory: In the procedural task (finger tapping), amount of entered sequences, amount of correctly entered sequences, and reaction time served as dependent measures and were each assessed by a 2 (time: pre vs post nap) x 2 (condition: stimulation vs sham) repeated measures ANOVA. Each ANOVA showed a significant main effect of time, demonstrating that participants entered more sequences after the nap (*M* = 22.93, *SE* = .71) than before (*M* = 20.58, *SE* = .70), completed more accurate sequences in the blocks after the nap (*M* = 20.94, *SE* = .78) than before (*M* = 18.46, *SE* = .69), and pressed buttons at a greater speed (pre: *M* = 247.40, *SE* = 13.17, post: *M* = 212.36, *SE* = 10.30; *F* (1,19) = 15.10, *p* < .001, *η_p_^2^* = .44; *F* (1,19) = 14.51, *p* = .001, *η_p_^2^* = .43, and *F* (1,19) = 11.93, *p* = .003, *η_p_^2^* = .39). Given that the task design did not include a wake control, it is unclear whether the improvement is due to sleep, time that has passed, or due to a natural improvement in the task over time. There was no significant main effect of stimulation nor an interaction effect in any of the three ANOVA (sequences: stim: *F* (1,19) = 1.41, *p* = .250, *η_p_^2^* = .07, interaction: *F* (1,19) < .01, *p* = .983, *η_p_^2^* < .01; accurate sequences: stim: *F* (1,19) = .67, *p* = .423, *η_p_^2^* = .03, interaction: *F* (1,19) < .01, *p* = .974, *η_p_^2^* < .01; RT: stim: *F* (1,19) = .86, *p* = .364, *η_p_^2^* = .04, interaction: *F* (1,19) = .14, *p* = .709, *η_p_^2^* = .01; Supplementary Fig. 4a-c). Some assumptions for running ANOVA were violated, so we separately also calculated the interaction effect by comparing the difference score in timing (pre-post) for each the stimulation and sham condition. Again, there was no significant effect of interaction this way for completed sequences (*V* = 88.5, *p* = .550), accurate sequences (*V* = 86.5, *p* = .502), or RT (*t*(19) = .38, *p* = .709, *d* = .08).

### Cortical spread of TSWs with swapped label-randomization

Statistical significance for differences in cortical spread of TSWs between the stimulation and sham condition was additionally assessed by randomly swapping condition labels in participants. Observed values were compared with a random distribution created by randomly swapped condition labels (1000 times). This procedure was used both for the count of each sensor pair involved in TSWs (Supplementary Fig. 5 a) and for the cortical spread of waves, corrected by total TSWs measured (Supplementary Fig. 5 b).

---

**Supplementary Table 1.**
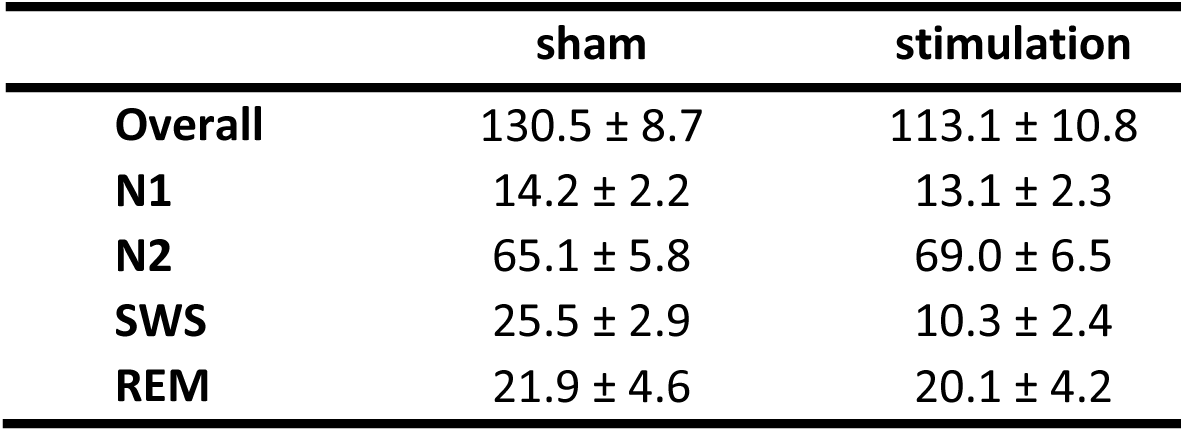
Sleep parameters for stimulation and sham condition. Duration is shown in minutes, by mean and standard error of the mean.

**Supplementary Figure 1.**
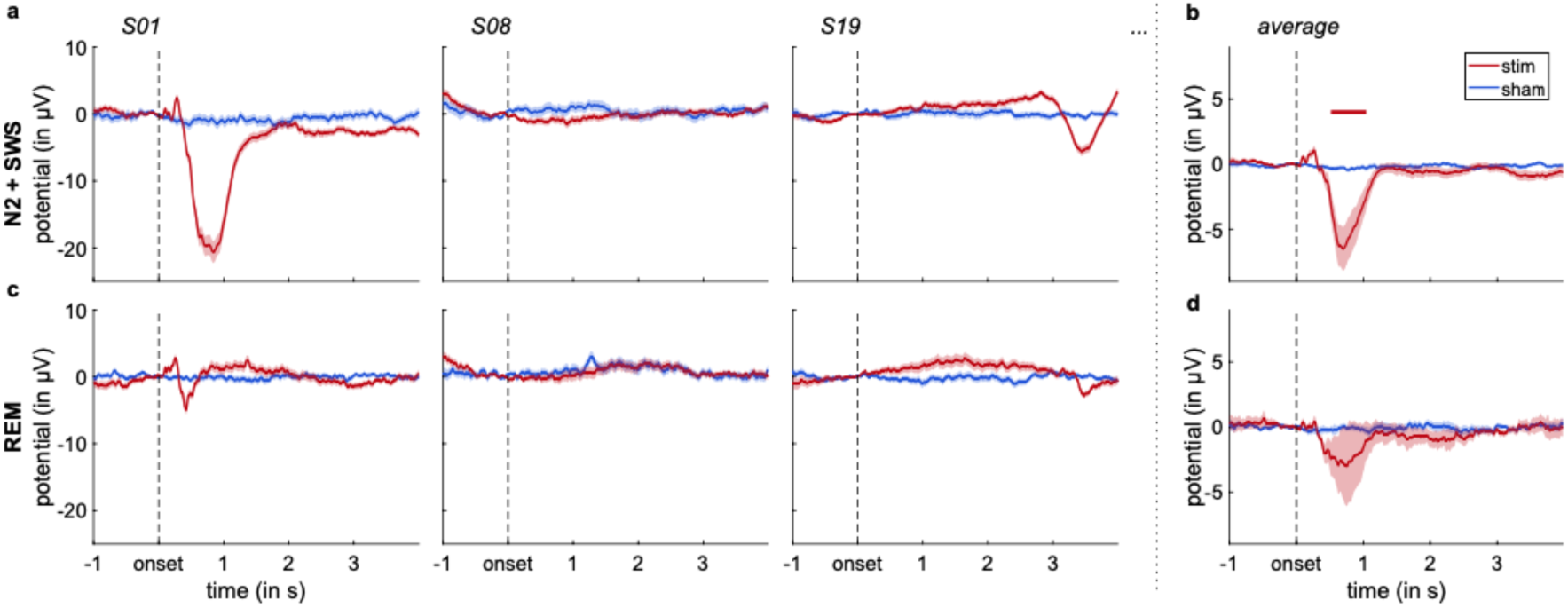
Auditory evoked responses in NREM and REM sleep. **a.** Event-related potentials for NREM sleep (N2 and SWS) in response to stimulation plotted for three individual exemplary subjects, one each for immediate (n=11), no (n=7), and delayed evoked potentials (n=2). ERPs show k complex-like inflections in the stimulation condition (red) in contrast to the sham condition (blue). Lines denote trial averages, shaded area standard error. **b.** Average ERP for NREM sleep. Red bar denotes time of significant difference across conditions. **c.** ERPs for REM sleep in response to stimulation plotted for three example subjects. **d.** Average ERP for REM sleep.

**Supplementary Figure 2.**
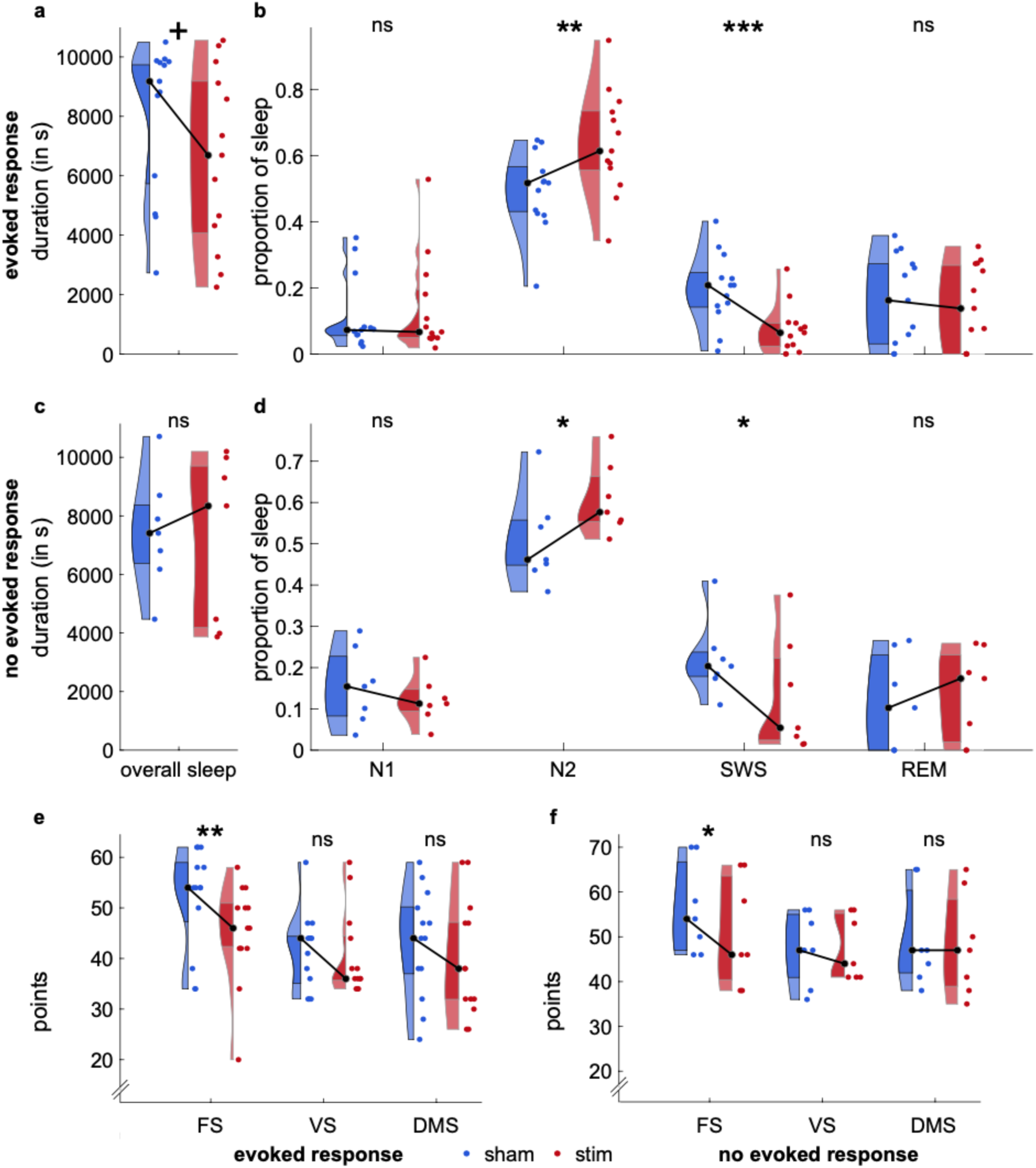
Effects of auditory stimulation in participants with and without auditory evoked responses in electrical brain activity. **a.** Sleep duration (in seconds) in sham (shown in blue) and stim conditions (shown in red) for subjects with evoked responses to stimulation. b. Proportion of time spent in each sleep stage, relative to the total duration, subjects as in a. Stimulation significantly reduced SWS and increased N2 proportion. c. Sleep duration (in seconds) in sham (shown in blue) and stim conditions (shown in red) for subjects without evoked responses to stimulation. d. Proportion of time spent in each sleep stage, relative to the total duration, subjects as in b. Stimulation significantly reduced SWS and increased N2 proportion. e. Declarative memory performance, shown as figural, verbal, and overall declarative memory score after the stimulation (red) and non-stimulation nap (blue), subjects as in a. Stimulation significantly reduced the figural memory score. f. Declarative memory performance, shown as figural, verbal, and overall declarative memory score after the stimulation (red) and non-stimulation nap (blue), subjects as in b. Stimulation significantly reduced the figural memory score.

**Supplementary Figure 3.**
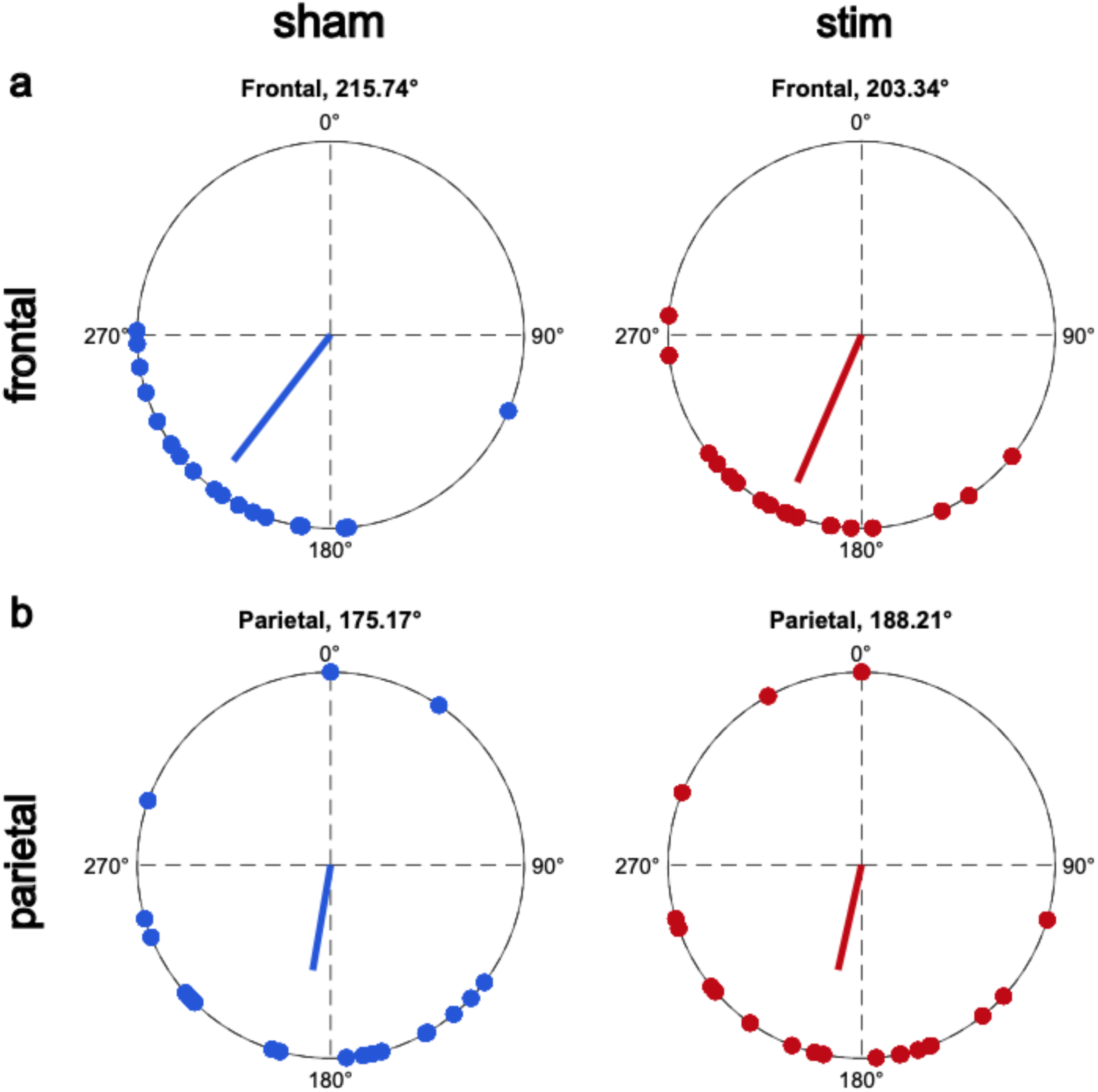
Phase coupling of slow waves and sleep spindles. **a.** Mean distribution of slow wave phase in which spindles originated in frontal channels (Fz, F3, F4) across subjects. No difference of angular mean phase was observed for stimulation (red) as opposed to sham (blue) condition. **b.** Same as a. but in parietal channels (Pz, P3, P4). No difference in angular mean phase was observed across conditions.

**Supplementary Figure 4.**
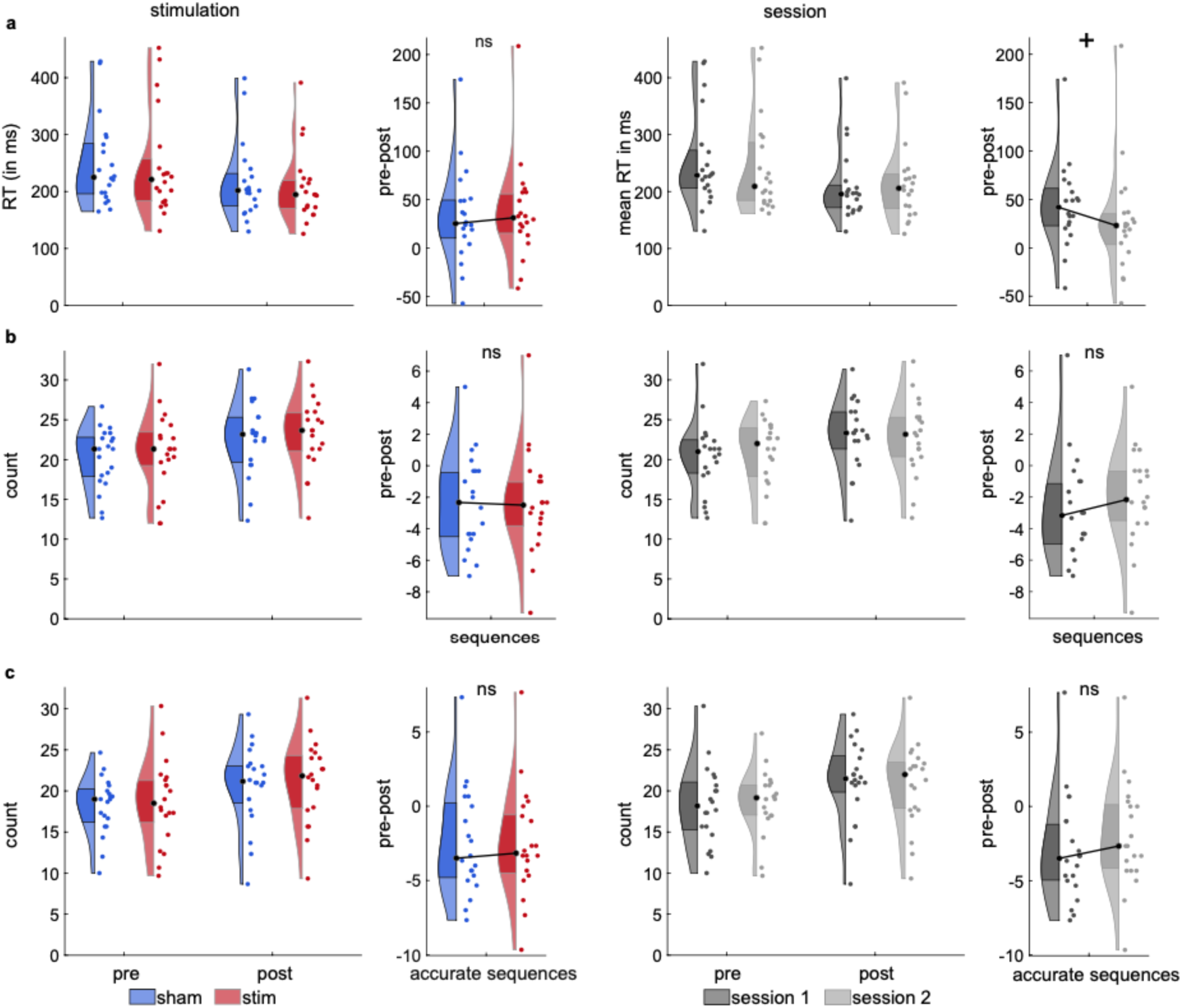
No effects of auditory stimulation on motor sequence memory. **a.** Speed (in ms) of fingertapping is displayed for the encoding part (pre nap) and the retrieval part (post nap), as well as the difference score with larger values indicating improvement. Colored left-hand plots show these differences separated by stimulation condition (sham in blue, stimulation in red), and right-hand plots separated by session (first in dark gray, second in light gray). Overall faster reaction times were observed post rather than pre nap. No significant condition differences occurred due to stimulation, but participants showed larger speed improvement in the first experimental session than in the second. **b.** Same as in a. but indicating how many sequences were typed out. Difference scores did not differ based on stimulation or session. **c.** Same as in a. but indicating how many accurate sequences were typed out. Difference scores did not differ based on stimulation or session.

**Supplementary Figure 5.**
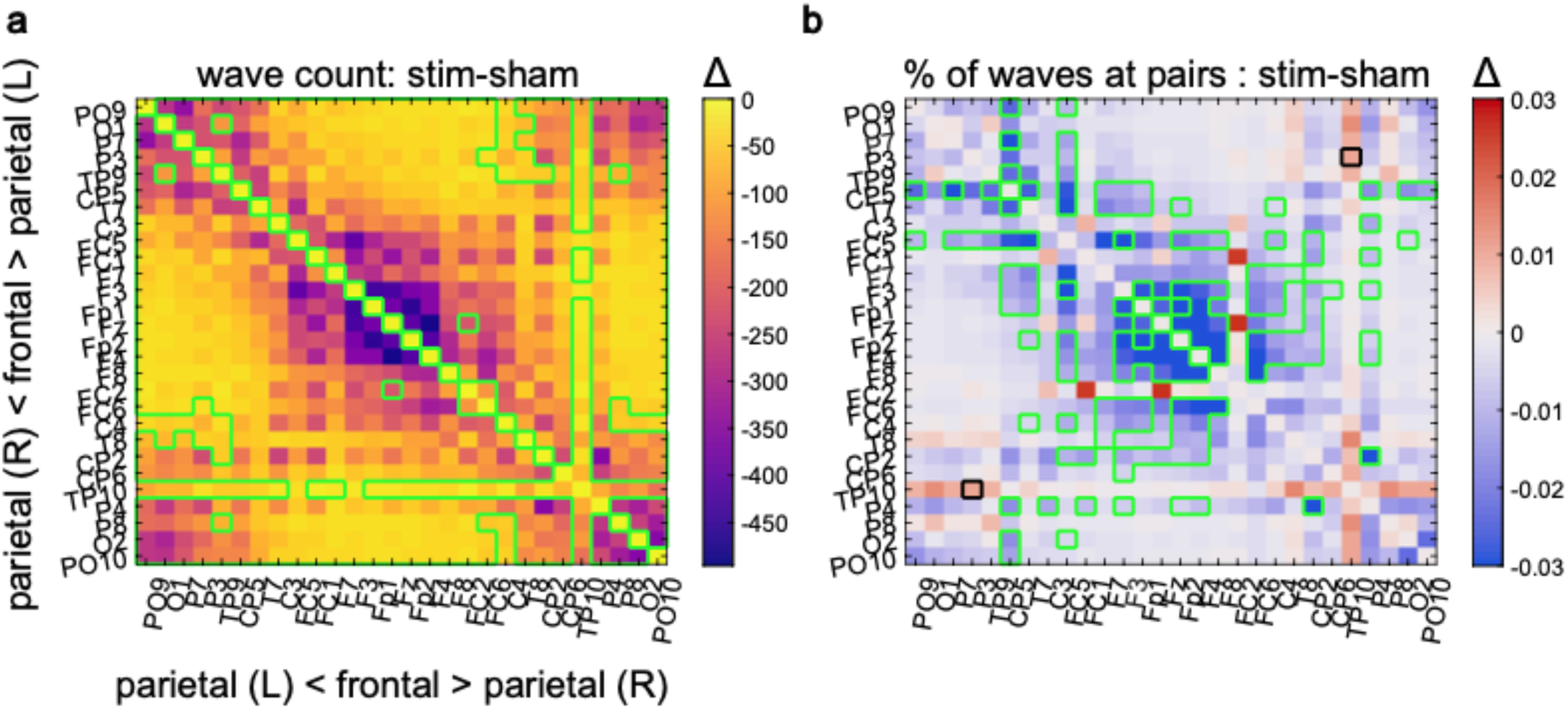
Effects of auditory stimulation on cortical spread of TSWs, swapped condition label randomization. **a.** The condition difference is shown between the count of TSWs involving each sensor pair and illustrates the scalp-wide reduction in traveling waves in the stimulation condition. Data as in Fig. 5a, significance indicators by swapped-label randomization and mirrored along the diagonal axis. Purple color indicates larger differences, thereby relatively higher TSW count in the sham condition. **b.** Difference in cortical spread of TSWs in each condition (as percent of TSWs involving each sensor pair) indicated broadly more scalp coverage for TSWs in sham (in blue), particularly in frontal sensors. Data as in Fig. 5i, significance indicators by swapped-label randomization and mirrored along the diagonal axis. Blue tones denote more relative TSW engagement of the region in the sham condition, red more relative TSW engagement in the stim condition. Green signifies the observed value below 2.5% of the label-flipped distribution, black above 97.5%, in line with an alpha of .05. Sensors are arranged from right parietal over frontal to left parietal.

## Notes

### Competing Interest Statement

The authors have declared no competing interest.

## References

Andrillon, T., Nir, Y., Staba, R. J., Ferrarelli, F., Cirelli, C., Tononi, G., & Fried, I. (2011). Sleep Spindles in Humans: Insights from Intracranial EEG and Unit Recordings. The Journal of Neuroscience, 31(49), 17821–17834.

Avvenuti, G., Handjaras, G., Betta, M., Cataldi, J., Imperatori, L. S., Lattanzi, S., Riedner, B. A., Pietrini, P., Ricciardi, E., Tononi, G., Siclari, F., Polonara, G., Fabri, M., Silvestrini, M., Bellesi, M., & Bernardi, G. (2020). Integrity of Corpus Callosum Is Essential for the Cross-Hemispheric Propagation of Sleep Slow Waves: A High-Density EEG Study in Split-Brain Patients. The Journal of Neuroscience, 40(29), 5589–5603. 10.1523/JNEUROSCI.2571-19.2020

Backhaus, J., Born, J., Hoeckesfeld, R., Fokuhl, S., Hohagen, F., & Junghanns, K. (2007). Midlife decline in declarative memory consolidation is correlated with a decline in slow wave sleep. Learning & Memory, 14(5), 336–341. 10.1101/lm.470507

Bakdash, J. Z., & Marusich, L. R. (2017). Repeated Measures Correlation. Frontiers in Psychology, 8, 456. 10.3389/fpsyg.2017.00456

Bäumler, G. (1974). Lern-und Gedächtnistest LGT-3. Hogrefe.

Belal, S., Cousins, J., El-Deredy, W., Parkes, L., Schneider, J., Tsujimura, H., Zoumpoulaki, A., Perapoch, M., Santamaria, L., & Lewis, P. (2018). Identification of memory reactivation during sleep by EEG classification. NeuroImage, 176, 203–214. 10.1016/j.neuroimage.2018.04.029

Bergmann, T. O., Mölle, M., Schmidt, M. A., Lindner, C., Marshall, L., Born, J., & Siebner, H. R. (2012). EEG-guided transcranial magnetic stimulation reveals rapid shifts in motor cortical excitability during the human sleep slow oscillation. Journal of Neuroscience, 32(1), 243–253. 10.1523/JNEUROSCI.4792-11.2012

Bernardi, G., Avvenuti, G., Cataldi, J., Lattanzi, S., Ricciardi, E., Polonara, G., Silvestrini, M., Siclari, F., Fabri, M., & Bellesi, M. (2021). Role of corpus callosum in sleep spindle synchronization and coupling with slow waves. Brain Communications, 3(2), fcab108. 10.1093/braincomms/fcab108

Bernardi, G., Siclari, F., Handjaras, G., Riedner, B. A., & Tononi, G. (2018). Local and Widespread Slow Waves in Stable NREM Sleep: Evidence for Distinct Regulation Mechanisms. Frontiers in Human Neuroscience, 12, 248. 10.3389/fnhum.2018.00248

Besedovsky, L., Ngo, H.-V. V., Dimitrov, S., Gassenmaier, C., Lehmann, R., & Born, J. (2017). Auditory closed-loop stimulation of EEG slow oscillations strengthens sleep and signs of its immune-supportive function. Nature Communications, 8(1984), 1–8. 10.1038/s41467-017-02170-3

Bódizs, R., Horváth, C. G., Szalárdy, O., Ujma, P. P., Simor, P., Gombos, F., Kovács, I., Genzel, L., & Dresler, M. (2022). Sleep-spindle frequency: Overnight dynamics, afternoon nap effects, and possible circadian modulation. Journal of Sleep Research, 31(3), e13514. 10.1111/jsr.13514

Born, J., & Wilhelm, I. (2012). System consolidation of memory during sleep. Psychological Research, 76(2), 192–203. 10.1007/s00426-011-0335-6

Cairney, S. A., Guttesen, A. á.Váli, El Marj, N., & Staresina, B. P. (2018). Memory Consolidation Is Linked to Spindle-Mediated Information Processing during Sleep. Current Biology, 28(6), 948–954.e4. 10.1016/j.cub.2018.01.087

Castelnovo, A., Lividini, A., Riedner, B. A., Avvenuti, G., Jones, S. G., Miano, S., Tononi, G., Manconi, M., & Bernardi, G. (2023). Origin, synchronization, and propagation of sleep slow waves in children. NeuroImage, 274, 120133. 10.1016/j.neuroimage.2023.120133

Cordi, M. J., & Rasch, B. (2021). No evidence for intra-individual correlations between sleep-mediated declarative memory consolidation and slow-wave sleep. SLEEP, 44(8), zsab034. 10.1093/sleep/zsab034

Cox, R., Schapiro, A. C., Manoach, D. S., & Stickgold, R. (2017). Individual Differences in Frequency and Topography of Slow and Fast Sleep Spindles. Frontiers in Human Neuroscience, 11, 433. 10.3389/fnhum.2017.00433

Das, A., Zabeh, E., Ermentrout, B., & Jacobs, J. (2024). Planar, Spiral, and Concentric Traveling Waves Distinguish Cognitive States in Human Memory. 10.1101/2024.01.26.577456

Dickey, C. W., Sargsyan, A., Madsen, J. R., Eskandar, E. N., Cash, S. S., & Halgren, E. (2021). Travelling spindles create necessary conditions for spike-timing-dependent plasticity in humans. Nature Communications, 12(1027), 1–15. 10.1038/s41467-021-21298-x

Diekelmann, S., & Born, J. (2010). The memory function of sleep. Nature Reviews, 11(2), 114–126. 10.1038/nrn2762

Dinges, D. F., & Powell, J. W. (1985). Microcomputer analyses of performance on a portable, simple visual RT task during sustained operations. Behavior Research Methods, Instruments, & Computers, 17(6), 652–655.

Drummond, S. P., Bischoff-Grethe, A., Dinges, D. F., Ayalon, L., Mednick, S. C., & Meloy, M. (2005). The neural basis of the psychomotor vigilance task. Sleep, 28(9), 1059–1068.

Dupret, D., O’Neill, J., Pleydell-Bouverie, B., & Csicsvari, J. (2010). The reorganization and reactivation of hippocampal maps predict spatial memory performance. Nature Neuroscience, 13(8), 995–1002. 10.1038/nn.2599

Esfahani, M. J., Farboud, S., Ngo, H.-V. V., Schneider, J., Weber, F. D., Talamini, L. M., & Dresler, M. (2023). Closed-loop auditory stimulation of sleep slow oscillations: Basic principles and best practices. Neuroscience & Biobehavioral Reviews, 153(September), 105379. 10.1016/j.neubiorev.2023.105379

Fehér, K. D., Omlin, X., Tarokh, L., Schneider, C. L., Morishima, Y., Züst, M. A., Wunderlin, M., Koenig, T., Hertenstein, E., Ellenberger, B., Ruch, S., Schmidig, F., Mikutta, C., Trinca, E., Senn, W., Feige, B., Klöppel, S., & Nissen, C. (2023). Feasibility, efficacy, and functional relevance of automated auditory closed-loop suppression of slow-wave sleep in humans. Journal of Sleep Research, 32(4), 1–13. 10.1111/jsr.13846

Fehér, K. D., Wunderlin, M., Maier, J. G., Hertenstein, E., Schneider, C. L., Mikutta, C., Züst, M. A., Klöppel, S., & Nissen, C. (2021). Shaping the slow waves of sleep: A systematic and integrative review of sleep slow wave modulation in humans using non-invasive brain stimulation. Sleep Medicine Reviews, 58(101438), 1–16. 10.1016/j.smrv.2021.101438

Feld, G. B., Wilhelm, I., Ma, Y., Groch, S., Binkofski, F., Mölle, M., & Born, J. (2013). Slow Wave Sleep Induced by GABA Agonist Tiagabine Fails to Benefit Memory Consolidation. Sleep, 36(9), 1317–1326. 10.5665/sleep.2954

Furuya, S., & Altenmüller, E. (2013). Flexibility of movement organization in piano performance. Frontiers in Human Neuroscience, 7. 10.3389/fnhum.2013.00173

Gais, S., Mölle, M., Helms, K., & Born, J. (2002). Learning-Dependent Increases in Sleep Spindle Density. The Journal of Neuroscience, 22(15), 6830–6834. 10.1523/JNEUROSCI.22-15-06830.2002

Halász, P. (1993). Arousals without awakening—Dynamic aspect of sleep. Physiology & Behavior, 54(4), 795–802. 10.1016/0031-9384(93)90094-V

Halász, P. (2016). The K-complex as a special reactive sleep slow wave – A theoretical update. Sleep Medicine Reviews, 29, 34–40. 10.1016/j.smrv.2015.09.004

Hangya, B., Tihanyi, B. T., Entz, L., Fabo, D., Eróss, L., Wittner, L., Jakus, R., Varga, V., Freund, T. F., & Ulbert, I. (2011). Complex Propagation Patterns Characterize Human Cortical Activity during Slow-Wave Sleep. Journal of Neuroscience, 31(24), 8770–8779. 10.1523/JNEUROSCI.1498-11.2011

Henin, S., Borges, H., Shankar, A., Sarac, C., Melloni, L., Friedman, D., Flinker, A., Parra, L. C., Buzsaki, G., Devinsky, O., & Liu, A. (2019). Closed-Loop Acoustic Stimulation Enhances Sleep Oscillations But Not Memory Performance. Eneuro, 6(6), ENEURO.0306-19.2019. 10.1523/ENEURO.0306-19.2019

Hu, X., Cheng, L. Y., Chiu, M. H., & Paller, K. A. (2020). Promoting memory consolidation during sleep: A meta-analysis of targeted memory reactivation. Psychological Bulletin, 146(3), 218–244. 10.1037/bul0000223

Kaestner, E. J., Wixted, J. T., & Mednick, S. C. (2013). Pharmacologically Increasing Sleep Spindles Enhances Recognition for Negative and High-arousal Memories. Journal of Cognitive Neuroscience, 25(10), 1597–1610. 10.1162/jocn_a_00433

Klinzing, J. G., Niethard, N., & Born, J. (2019). Mechanisms of systems memory consolidation during sleep. Nature Neuroscience, 22, 1596–1610. 10.1038/s41593-019-0467-3

Klinzing, J. G., Tashiro, L., Ruf, S., Wolff, M., Born, J., & Ngo, H.-V. V. (2021). Auditory stimulation during sleep suppresses spike activity in benign epilepsy with centrotemporal spikes. Cell Reports Medicine, 2(11), 100432. 10.1016/j.xcrm.2021.100432

Kokkinos, V., & Kostopoulos, G. K. (2011). Human non-rapid eye movement stage II sleep spindles are blocked upon spontaneous K-complex coincidence and resume as higher frequency spindles afterwards. Journal of Sleep Research, 20(1pt1), 57–72. 10.1111/j.1365-2869.2010.00830.x

Kumral, D., Matzerath, A., Leonhart, R., & Schönauer, M. (2023). Spindle-dependent memory consolidation in healthy adults: A meta-analysis. Neuropsychologia, 189, 108661. 10.1016/j.neuropsychologia.2023.108661

Kurth, S., Riedner, B. A., Dean, D. C., O’Muircheartaigh, J., Huber, R., Jenni, O. G., Deoni, S. C. L., & LeBourgeois, M. K. (2017). Traveling Slow Oscillations During Sleep: A Marker of Brain Connectivity in Childhood. Sleep, 40(9). 10.1093/sleep/zsx121

Latchoumane, C.-F. V., Ngo, H. V., Born, J., & Shin, H.-S. (2017). Thalamic Spindles Promote Memory Formation during Sleep through Triple Phase-Locking of Cortical, Thalamic, and Hippocampal Rhythms. Neuron, 95(2), 424–435. 10.1016/j.neuron.2017.06.025

Lechat, B., Hansen, K., Micic, G., Decup, F., Dunbar, C., Liebich, T., Catcheside, P., & Zajamsek, B. (2021). K-complexes are a sensitive marker of noise-related sensory processing during sleep: A pilot study. Sleep, 44(9), zsab065. 10.1093/sleep/zsab065

Lubenov, E. V., & Siapas, A. G. (2009). Hippocampal theta oscillations are travelling waves. Nature, 459(7246), 534–539. 10.1038/nature08010

Mander, B. A., Marks, S. M., Vogel, J. W., Rao, V., Lu, B., Saletin, J. M., Ancoli-Israel, S., Jagust, W. J., & Walker, M. P. (2015). β-amyloid disrupts human NREM slow waves and related hippocampus-dependent memory consolidation. Nature Neuroscience, 18(7), 1051–1057. 10.1038/nn.4035

Manoach, D. S., Thakkar, K. N., Stroynowski, E., Ely, A., McKinley, S. K., Wamsley, E., Djonlagic, I., Vangel, M. G., Goff, D. C., & Stickgold, R. (2010). Reduced overnight consolidation of procedural learning in chronic medicated schizophrenia is related to specific sleep stages. Journal of Psychiatric Research, 44(2), 112–120. 10.1016/j.jpsychires.2009.06.011

Marshall, L., Helgadóttir, H., Mölle, M., & Born, J. (2006). Boosting slow oscillations during sleep potentiates memory. Nature, 444(7119), 610–613. 10.1038/nature05278

Massimini, M., Ferrarelli, F., Esser, S. K., Riedner, B. A., Huber, R., Murphy, M., Peterson, M. J., & Tononi, G. (2007). Triggering sleep slow waves by transcranial magnetic stimulation. Proceedings of the National Academy of Sciences of the United States of America, 104(20), 8496–8501. 10.1073/pnas.0702495104

Massimini, M., Huber, R., Ferrarelli, F., Hill, S., & Tononi, G. (2004). The Sleep Slow Oscillation as a Traveling Wave. The Journal of Neuroscience, 24(31), 6862–6870. 10.1523/JNEUROSCI.1318-04.2004

Muller, L., Chavane, F., Reynolds, J., & Sejnowski, T. J. (2018). Cortical travelling waves: Mechanisms and computational principles. Nature Reviews Neuroscience, 19(5), 255–268. 10.1038/nrn.2018.20

Muller, L., Piantoni, G., Koller, D., Cash, S. S., Halgren, E., & Sejnowski, T. J. (2016). Rotating waves during human sleep spindles organize global patterns of activity that repeat precisely through the night. eLife, 5, e17267. 10.7554/eLife.17267

Muzet, A. (2007). Environmental noise, sleep and health. Sleep Medicine Reviews, 11(2), 135–142. 10.1016/j.smrv.2006.09.001

Ngo, H.-V., Fell, J., & Staresina, B. (2020). Sleep spindles mediate hippocampal-neocortical coupling during long-duration ripples. eLife, 9, e57011. 10.7554/eLife.57011

Ngo, H.-V. V., Martinetz, T., Born, J., & Mölle, M. (2013). Auditory Closed-Loop Stimulation of the Sleep Slow Oscillation Enhances Memory. Neuron, 78, 545–553. 10.1016/j.neuron.2013.03.006

Nishida, M., & Walker, M. P. (2007). Daytime naps, motor memory consolidation and regionally specific sleep spindles. PLoS ONE, 2(4), e341: 1-7. 10.1371/journal.pone.0000341

Oldfield, R. C. (1971). The assessment and analysis of handedness: The Edinburgh inventory. Neuropsychologia, 9(1), 97–113.

Oostenveld, R., Fries, P., Maris, E., & Schoffelen, J.-M. (2011). FieldTrip: Open Source Software for Advanced Analysis of MEG, EEG, and Invasive Electrophysiological Data. Computational Intelligence and Neuroscience, 2011, 1–9. 10.1155/2011/156869

Oudiette, D., & Paller, K. A. (2013). Upgrading the sleeping brain with targeted memory reactivation. Trends in Cognitive Sciences, 17(3), 142–149. 10.1016/j.tics.2013.01.006

Petzka, M., Chatburn, A., Charest, I., Balanos, G. M., & Staresina, B. P. (2022). Sleep spindles track cortical learning patterns for memory consolidation. Current Biology, 32(11), 2349–2356.e4. 10.1016/j.cub.2022.04.045

Plihal, W., & Born, J. (1997). Effects of Early and Late Nocturnal Sleep on Declarative and Procedural Memory. Journal of Cognitive Neuroscience, 9(4), 534–547. 10.1162/jocn.1997.9.4.534

Rasch, B., & Born, J. (2013). About Sleep’s Role in Memory. Physiological Review, 93, 681– 766. 10.1152/physrev.00032.2012

Rasch, B., Christian, Büchel., Gais, S., & Born, J. (2007). Odor Cues During Slow-Wave Sleep Prompt Declarative Memory Consolidation. Science, 315(5817), 1426–1429.

Rechtschaffen, A., & Kales, A. (1968). A Manual of Standardized Terminology, Techniques and Scoring System for Sleep Stages of Human Subjects. U.S. Department of Health, Education and Welfare Public Health Service - National Institutes of Health.

Roenneberg, T., Kuehnle, T., Juda, M., Kantermann, T., Allebrandt, K., Gordijn, M., & Merrow, M. (2007). Epidemiology of the human circadian clock. Sleep Medicine Reviews, 11(6), 429–438.

Rudoy, J. D., Voss, J. L., Westerberg, C. E., & Paller, K. A. (2009). Strengthening Individual Memories by Reactivating Them During Sleep. Science, 326(November), 1079. 10.1126/science.1179013

Schabus, M., Hödlmoser, K., Gruber, G., Sauter, C., Anderer, P., Klösch, G., Parapatics, S., Saletu, B., Klimesch, W., & Zeitlhofer, J. (2006). Sleep spindle-related activity in the human EEG and its relation to general cognitive and learning abilities. European Journal of Neuroscience, 23, 1738–1746. 10.1111/j.1460-9568.2006.04694.x

Schoch, S. F., Cordi, M. J., Schredl, M., & Rasch, B. (2018). The effect of dream report collection and dream incorporation on memory consolidation during sleep.

Schönauer, M., Alizadeh, S., Jamalabadi, H., Abraham, A., Pawlizki, A., & Gais, S. (2017). Decoding material-specific memory reprocessing during sleep in humans. Nature Communications, 8(15404), 1–9. 10.1038/ncomms15404

Schönauer, M., Geisler, T., & Gais, S. (2013). Strengthening Procedural Memories by Reactivation in Sleep. Journal of Cognitive Neuroscience, 26(1), 143–153. 10.1162/jocn

Schönauer, M., Pawlizki, A., Köck, C., & Gais, S. (2014). Exploring the effect of sleep and reduced interference on different forms of declarative memory. Sleep, 37(12), 1995– 2007. 10.5665/sleep.4258

Schreiner, T., Griffiths, B. J., Kutlu, M., Vollmar, C., Kaufmann, E., Quach, S., Remi, J., Noachtar, S., & Staudigl, T. (2024). Spindle-locked ripples mediate memory reactivation during human NREM sleep. Nature Communications, 15(1), 5249. 10.1038/s41467-024-49572-8

Schreiner, T., Petzka, M., Staudigl, T., & Staresina, B. P. (2023). Respiration modulates sleep oscillations and memory reactivation in humans. Nature Communications, 14(1), 8351. 10.1038/s41467-023-43450-5

Schreiner, T., & Staudigl, T. (2020). Electrophysiological signatures of memory reactivation in humans. Philosophical Transactions of the Royal Society B, 375(20190293), 1–15.

Sharon, O., Chen, X., Dude, J., Shah, V. D., Ju, Y.-E. S., Jagust, W. J., & Walker, M. P. (2024). Tau pathology leads to lonely non-traveling slow waves that mediate human memory impairment. 10.1101/2024.05.22.595043

Siclari, F., Bernardi, G., Riedner, B. A., LaRocque, J. J., Benca, R. M., & Tononi, G. (2014). Two Distinct Synchronization Processes in the Transition to Sleep: A High-Density Electroencephalographic Study. Sleep, 37(10), 1621–1637. 10.5665/sleep.4070

Sousouri, G., Krugliakova, E., Skorucak, J., Leach, S., Snipes, S., Ferster, M. L., Da Poian, G., Karlen, W., & Huber, R. (2022). Neuromodulation by means of phase-locked auditory stimulation affects key marker of excitability and connectivity during sleep. Sleep, 45(1), zsab204. 10.1093/sleep/zsab204

Staresina, B. P., Bergmann, T. O., Bonnefond, M., Van Der Meij, R., Jensen, O., Deuker, L., Elger, C. E., Axmacher, N., & Fell, J. (2015). Hierarchical nesting of slow oscillations, spindles and ripples in the human hippocampus during sleep. Nature Neuroscience, 18(11), 1679–1686. 10.1038/nn.4119

Stickgold, R., & Walker, M. P. (2005). Memory consolidation and reconsolidation: What is the role of sleep? Trends in Neurosciences, 28(8), 408–415. 10.1016/j.tins.2005.06.004

Walker, M. P., Brakefield, T., Morgan, A., Hobson, J. A., & Stickgold, R. (2002). Practice with Sleep Makes Perfect: Sleep-Dependent Motor Skill Learning. Neuron, 35(1), 205–211.

Walker, M. P., Brakefield, T., Seidman, J., Morgan, A., Hobson, J. A., & Stickgold, R. (2003). Sleep and the Time Course of Motor Skill Learning. Learning & Memory, 10(4), 275–284. 10.1101/lm.58503

Walker, M. P., Stickgold, R., Alsop, D., Gaab, N., & Schlaug, G. (2005). Sleep-dependent motor memory plasticity in the human brain. Neuroscience, 133(4), 911–917. 10.1016/j.neuroscience.2005.04.007

Wamsley, E. J., Tucker, M. A., Shinn, A. K., Ono, K. E., McKinley, S. K., Ely, A. V., Goff, D. C., Stickgold, R., & Manoach, D. S. (2012). Reduced Sleep Spindles and Spindle Coherence in Schizophrenia: Mechanisms of Impaired Memory Consolidation? Biological Psychiatry, 71(2), 154–161. 10.1016/j.biopsych.2011.08.008

Westerberg, C. E., Florczak, S. M., Weintraub, S., Mesulam, M., Marshall, L., Zee, P. C., & Paller, K. A. (2015). Memory improvement via slow-oscillatory stimulation during sleep in older adults. Neurobiology of Aging, 36(9), 2577–2586. 10.1016/j.neurobiolaging.2015.05.014

Wilckens, K. A., Ferrarelli, F., Walker, M. P., & Buysse, D. J. (2018). Slow-Wave Activity Enhancement to Improve Cognition. Trends in Neurosciences, 41(7), 470–482. 10.1016/j.tins.2018.03.003

Wunderlin, M., Züst, M. A., Hertenstein, E., Fehér, K. D., Schneider, C. L., Klöppel, S., & Nissen, C. (2021). Modulating overnight memory consolidation by acoustic stimulation during slow-wave sleep: A systematic review and meta-analysis. Sleep, 44(7), zsaa296. 10.1093/sleep/zsaa296

Zhang, H., Fell, J., & Axmacher, N. (2018). Electrophysiological mechanisms of human memory consolidation. Nature Communications, 9(4103), 1–11. 10.1038/s41467-018-06553-y

Zhang, H., & Jacobs, J. (2015). Traveling Theta Waves in the Human Hippocampus. 35(36), 12477–12487. 10.1523/JNEUROSCI.5102-14.2015

